# An aiHumanoid Simulation of Gram-Negative Sepsis: A Comprehensive Multi-Organoid Platform for Advanced Disease Modeling, Drug Discovery, and Personalized Medicine

**DOI:** 10.1101/2023.07.18.549304

**Authors:** Wayne R Danter

## Abstract

Organoids are three-dimensional cellular structures resembling human organs, which have emerged as valuable tools for studying organ development, disease modeling, and drug discovery. Integrating multiple organoid systems allows for the examination of complex interactions between different organs. In this study, we present the development and initial validation of the aiHumanoid simulation, an advanced AI-based computational framework that integrates 18 individual organoid simulations through a common cardiovascular system. Our aim is to investigate the systemic effects of gram-negative sepsis, a life-threatening condition that affects multiple organ systems.

In this study, we evaluated the impact of gram-negative sepsis on the organoid systems that make up the aiHumanoid simulation. Our findings indicate significant alterations in cardiovascular, nervous system, respiratory, renal, hepatic, hematologic, gastrointestinal, musculoskeletal, immune, and endocrine parameters in both male and female sepsis-affected organoids. Notably, markers of inflammation, coagulation, renal dysfunction, liver damage, immune response, and endocrine regulation were significantly affected by sepsis. While some parameters showed gender-specific differences in response, such as hormonal changes, the overall impact of gram-negative sepsis was observed in both sexes.

This study demonstrates the potential of the aiHumanoid to accurately simulate the systemic effects of diseases on various organ systems. The integration of computational simulations with organoid systems offers a powerful approach for understanding disease mechanisms and evaluating potential therapies. By providing a more efficient and physiologically relevant platform for drug testing, the aiHumanoid simulation has the potential to accelerate the drug development process, reduce costs, and minimize the need for animal testing. Further research and ongoing validation will be crucial to fully exploit the capabilities of this revolutionary computational framework for advancing disease modeling and therapeutic interventions.

## Introduction

Organoids are three-dimensional, self-organizing cellular structures that resemble the architecture and function of various organs in the human body [1]. These miniaturized and simplified versions of organs are derived from pluripotent stem cells or organ-specific adult stem cells, which give rise to different cell types present in the organ they mimic [2]. Organoids have emerged as a valuable in vitro model for studying organ development, tissue homeostasis, disease modeling, and drug discovery [3,4].

The integration of multiple organoid systems presents an opportunity to study the complex interplay between different organs within the human body. Combining organoids enables researchers to better understand the systemic effects of diseases and the response to therapeutic interventions [5]. The development of interconnected organoid systems has the potential to advance our understanding of human biology and provide a more accurate representation of the physiological and pathological processes occurring in the human body.

In recent years, computer simulations of organoids have been developed to complement and enhance experimental studies. These computational models provide a powerful tool for predicting organoid behavior, testing hypotheses, and identifying optimal conditions for organoid growth and differentiation [6]. Hofer and Lutolf discuss the engineering of organoids and their potential uses in biomedical research [16]. Furthermore, combining computer simulations of multiple organoid systems can facilitate the study of complex diseases, such as sepsis, which involve the interaction of multiple organs [7].

Developing the aiHumanoid simulation with integrated organoid systems poses several technical challenges, such as creating a standardized simulation platform, ensuring organoid compatibility, and calibrating organoid models [8,9]. Addressing these challenges is critical to constructing a reliable and accurate computational framework that can effectively mimic the complex interplay between different organ systems. In this work, we detail the methods employed to overcome these challenges and validate the aiHumanoid simulation.

A validated aiHumanoid simulation has the potential to revolutionize disease modeling, drug discovery, and repurposing by providing a more accurate and efficient in silico platform for testing potential therapies [10]. The simulation could reduce the need for in vitro and in vivo laboratory testing and expedite the drug development process, enabling a more seamless transition to Phase 1 human trials [11]. This computational approach could reduce the time and cost associated with drug development while minimizing the use of animal testing and providing more physiologically relevant data [12,13].

In this paper, we extend the research reporting on the development of aiWBO and aiLUNG (whole brain organoids and lung organoids) published previously by Dr Sally Ezra and me [14,15] Here, we present the development and initial validation of the aiHumanoid simulation, an advanced AI based computational framework that integrates 18 individual organoid simulations through a common cardiovascular system (CVS). This aiHumanoid simulation aims to provide a platform for investigating the systemic effects of diseases and the response to therapeutic interventions. We demonstrate the utility of our model by simulating gram-negative sepsis (Pseudomonas Aeruginosa), a severe and life-threatening condition that affects all organ systems in the body.

The paper is divided in to two main parts. Part 1 deals with the creation and validation of the aiHumanoid simulations for both males and females. In Part 2 we evaluate the effects of a serious multiorgan disease, namely gram-negative sepsis on the male and female aiHumanoid simulations.

## Methods

The methodology for the generation and evaluation of artificial intelligence (AI) models of aiHumanoid begins with the simulation of induced pluripotent stem cells (aiPSCs). This approach is grounded in the pioneering research conducted by Yamanaka and colleagues [17]. The primary steps involve the construction of signal transduction pathways to emulate a fibroblast. Subsequently, this simulated entity is exposed to the quartet of Yamanaka transcription factors, specifically cMYC (cellular Myelocytomatosis proto-oncogene), OCT4 (octamer-binding transcription factor 4), KLF4 (Kruppel-like factor 4), and SOX2 ((sex determining region Y)-box 2), within a simulated growth and developmental medium. Comprehensive procedures for the creation and validation of aiPSCs, derived from simulated human fibroblasts, are documented in the initial DeepNEU technology overview [18].

Growth media development for simulating 18 individual organoid types presented a significant challenge. A commonality amongst many organoids is their use of growth media containing elements such as DMEM/F12 [19], with or without B27 supplement [20] and Matrigel [21]. After several trials, it became apparent that a unified growth medium for all organoid types might be feasible.

Growth media development began by identifying and comparing individual growth media for each organoid type. Shared characteristics were determined amongst them, leading to the formulation of a basic medium composed of DMEM/F12, B27 supplement, and Matrigel.

Subsequently, organoid-specific media modifications were made by removing the base ingredients-DMEM/F12, B27 supplement, and Matrigel, leaving only the elements unique to each organoid. This process necessitated the development of distinct medium recipes for each organoid type.

After considerable experimentation, specific media for bone marrow, cardiac, kidney, liver, lung, and thyroid organoids were successfully developed. The final cocktail included DMEM/F12, B27, Matrigel, bone marrow medium, cardiac medium, liver medium, kidney medium, lung medium, and thyroid medium. Additional organoid media for adrenal, brain stem, breast, cerebral, cerebellar, gallbladder, ovary, skin, and testis organoids were explored but did not improve on the specific organoid subset outlined above for aiHumanoid development.

While implementing this complex cocktail in a wet lab setting would be highly challenging, it proved to be a simple and effective solution for use in computer-simulated systems. This comprehensive growth medium now serves as the standard for all aiHumanoid simulations.

### 3.1. Creation of Wild-Type Organoid Simulations

After validation through peer-reviewed literature, artificial intelligence (AI) simulations of human induced pluripotent stem cells (aiPSC) were generated and utilized to derive various organoid types. These wild-type organoids were engineered to simulate a wide array of organoids and cellular types, emphasizing the importance of cellular diversity in ensuring the representation of expected cell types and expression patterns within the specific organoid type.

The comprehensive protocols used for generating lung and whole-brain organoids have been published [22, 23]. In broad terms, individual wild-type organoids were created through one of two pathways. The first, an unguided pathway, capitalizes on the tendency of pluripotent stem cells to differentiate along neural pathways over time under generalized conditions [3]. This approach is generally applicable to cerebral, cerebellar, and other central nervous system (CNS) organoids. Conversely, the second approach employs a combination of specialized media and specific elements such as transcription factors and hormones, that guide stem cells towards specific lineages [2, 24]. This guided approach is typically required for most non-CNS organoid development. The development of the aiHumanoid necessitates a more complex combination of a common basic medium and more specific cell line/organoid media, as detailed above.

### 3.2. Integration of Individual Organoids into the aiHumanoid Simulation

In vitro organoids depend on diffusion from the surrounding medium to supply glucose, oxygen, and other nutritional requirements, and to remove toxic metabolites, often necessitating frequent medium replacement. This method frequently results in restricted growth and central necrosis as the organoid growth outpaces the capabilities of diffusion. These recognized limitations of in vitro organoids are predominantly due to the absence of cardiovascular, respiratory, and renal systems.

To construct a more physiologically accurate multi-organoid system, we have integrated all individual organoids with a central cardiovascular system. This is achieved by creating arterial pathways to all organoids and venous return pathways from all organoids, including lung and renal organoids. This design should produce a more physiological scenario in terms of nutrition, oxygenation, toxic metabolite removal, and aiHumanoid growth and stability.

### 3.3 Validation of Individual Organoids into the aiHumanoid Simulation

Validation of the aiHumanoid simulations is based on an extensive list of organoid related markers derived from the published literature (see Appendix A for details).

### 3.4 Statistical Analysis

Both male and female aiHumanoid simulations generated data limited to the mean activity or expression levels ranging between 0 and +1, and a sample size (n) of 15 for each marker within each individual organoid system. The 95% Confidence Interval (CI) was calculated using the formula 95%CI(X) = X ±1.96*sqrt((X*(1-X))/15) [25]. Standard Deviation (SD) information from the raw data were not available and we were therefor unable to use the T-test or Mann-Whitney u test. To compensate for the lack of SD information we used the two tailed Z test to statistically evaluate the differences in male v. female and uninfected (WT) v gram-negative sepsis data [26,27]. This approach was used in all data analysis in Part 1 and Part 2 of the paper.

### 3.5 The Null hypothesis

For this project, the Null Hypothesis (H0) states that there is no significant difference in the mean expression levels or activity of the markers within individual organoid systems between male and female aiHumanoid simulations (Part 1), and between uninfected (WT) and gram-negative sepsis simulations (Part 2), at a significance level of p<0.05. This means that if the p-value resulting from the two tailed Z-test is less than 0.05, the null hypothesis would be rejected, indicating a significant difference in the means.

## Results-Part 1

AI platform metrics: This version of the DeepNEU platform (v8.0, 2023) has several improvements compared with the most recent previous version (v7.2, 2023). First, v8 contains 6946 genotypic and phenotypic concepts connected by 65560 positive and negative concepts. Secondly, this means there are more than 9.4 incoming and outgoing relationships for each nonzero concept in the relationship matrix. Thirdly, we have created, and literature validated a total of 18 organoid simulations compared with the previous 2 organoids in v7.2, namely aiWBO and aiLUNG. Next, we implemented the important Gut-Brain axis which was not implemented in previous versions. Finally, we have improved the early stopping algorithm based on unseen peer reviewed results. The multiple networks consistently converge at between 50 and 60 iterations. Based on the previously unseen validation data the networks produced optimal results after 13-16 iterations. Beyond that overfitting becomes noticeable. Using a 3 valued moving average, the results after 15 iterations was used for all simulations in all experiments.

The purpose of Part 1 of this study, was to evaluate the ability of the aiHumanoid simulations to express recognized multiple organ system markers. These markers (see Appendix A) were used to compare the normal physiological status of male and female organoid simulations.

### 1. CVS

#### 1.1. Overview of the aiHumanoid Cardiovascular System Simulation (CVS) Data

In this study, we present the initial validation data for an aiHumanoid simulation that integrates 18 individual organoid simulations through a common cardiovascular system. The evaluation was performed by comparing the expression levels of various cardiac and vascular markers in male and female simulations. The markers were analyzed to evaluate the development of a simulated cardiovascular system (CVS).

#### 1.2. Cardiac Marker (N=5) Expression in the aiHumanoid Simulation

The cardiac markers evaluated included ACTC1 (Actin Alpha Cardiac Muscle 1), MYH6 (Myosin Heavy Chain 6), MYH7 (Myosin Heavy Chain 7), NKX2.5 (NK2 Homeobox 5) and TNNT2 (Troponin T2, Cardiac Type).

ACTC1, MYH6, and NKX2.5 show no significant difference (NS, p>0.05) between males and females, all with high expression levels (H). MYH7, however, shows a medium level of expression (M) in both genders, with a statistically significant difference favoring males (p<0.05). TNNT2, displays low levels of expression (L) in both genders with no significant difference. These results are summarized in Table 1a.

#### 1.3. Vascular Marker (N=5) Expression in the aiHumanoid Simulation

We also assessed the expression of vascular markers, including aSMA (Alpha Smooth Muscle Actin), CD31 (Cluster of Differentiation 31), NG2 (Neuron-Glial Antigen 2), VE-Cadherin (Vascular Endothelial Cadherin), and VWF (Von Willebrand Factor). aSMA shows a significant difference (p<0.01) between males and females, with low (L) expression in males and medium (M) expression in females. CD31, NG2, VE-Cadherin, and VWF all exhibit high expression levels (H) with no significant difference between males and females. These results are summarized in Table 1b.

**Table 1:**
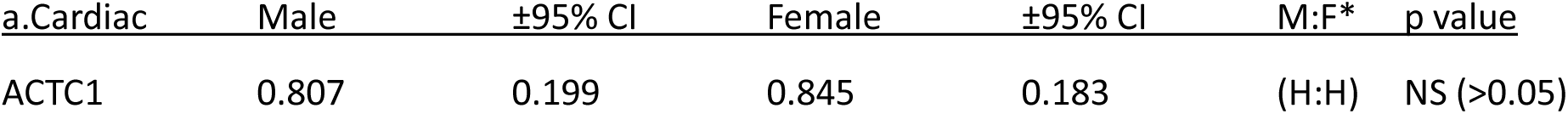
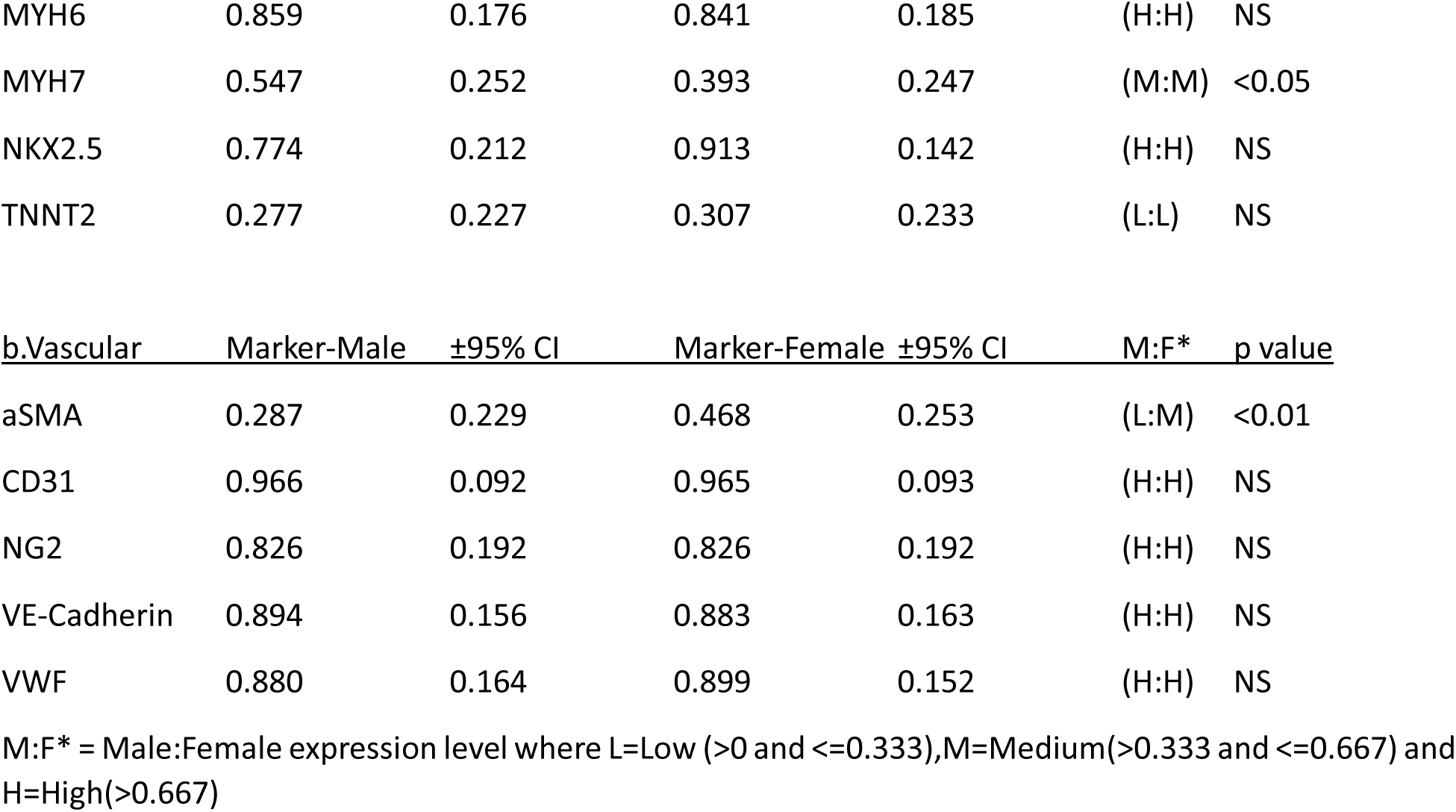
A comparison of male and female makers of a physiological CVS.

#### 1.4. Summary of the aiHumanoid Cardiovascular System Simulations

In summary, among the physiological CVS markers analyzed, MYH7 and aSMA showed significant gender differences, with MYH7 being more expressed in males, and aSMA more expressed in females. All other markers showed similar levels of expression between genders. These results indicate that the aiHumanoid simulations exhibit a consistent expression of cardiac and vascular markers across both male and female simulations. There appear to be only 2 statistically significant gender differences amongst 10 CVS markers.

### 2. CNS

#### 2.1. Overview of the aiHumanoid Central Nervous System (CNS) Simulations

Next, we examined the central nervous system (CNS) component of the aiHumanoid simulation by evaluating the cerebral, cerebellar and brain stem elements. A spinal cord simulation is in development for the next generation of simulations.

The expression levels of various regional markers, including astrocyte, oligodendrocyte, and neuronal markers were compared between male and female simulations to evaluate the performance of the simulated CNS.

#### 2.2. Cerebral Marker (N=5) Expression in the aiHumanoid Simulation

The cerebral markers analyzed in this study included CTIP2 (COUP-TF Interacting Protein 2), FOXG1 (Forkhead Box G1), PAX6 (Paired Box 6), SOX2 (SRY (Sex Determining Region Y)-Box 2), TBR1 (T-box Brain Protein 1). All cerebral markers were expressed in both genders. However, there were two statistically significant differences. The expression of CTIP2 was increased in females relative to males (p<0.01) while for TBR2 expression this gender difference was reversed (p<0.05) (in Table 2a).

#### 2.3. Astrocyte Marker (N=3) Expression in the aiHumanoid Simulation

We assessed the expression of astrocyte markers, including ALDH1L1 (Aldehyde Dehydrogenase 1 Family Member L1), GFAP (Glial Fibrillary Acidic Protein), and S100B (S100 Calcium Binding Protein B). All three markers were expressed in both genders and there were no significant differences identified (in Table 2b).

#### 2.4 Oligodendrocyte Marker (N=3) Expression in the aiHumanoid Simulation

The markers evaluated included MBP (Myelin Basic Protein), OLIG2 (Oligodendrocyte Transcription Factor 2), SOX10 (Sex Determining Region Y)-Box 10). All three markers were expressed in both genders and there were no significant differences identified (in Table 2c).

#### 2.5. Neuronal Marker (N=3) Expression in the aiHumanoid Simulation

The neuronal markers analyzed included MAP2 (Microtubule-Associated Protein 2), NeuN (Neuronal Nuclei), and TUBB3 (Tubulin Beta 3 Class III). MAP2 and TUBB3 were both expressed without significant gender differences. TUBB3 on the other hand demonstrated Low expression with significantly increased expression if female simulations (p<0.001) (in Table 2d).

#### 2.6. Evaluation of the aiHumanoid Central Nervous System Simulation

The results demonstrate that the aiHumanoid simulation consistently expresses cerebral, astrocyte, oligodendrocyte, and neuronal markers across both male and female simulations. This indicates that the aiHumanoid simulation serves as a promising platform for modeling the human central nervous system.

**Table 2:**
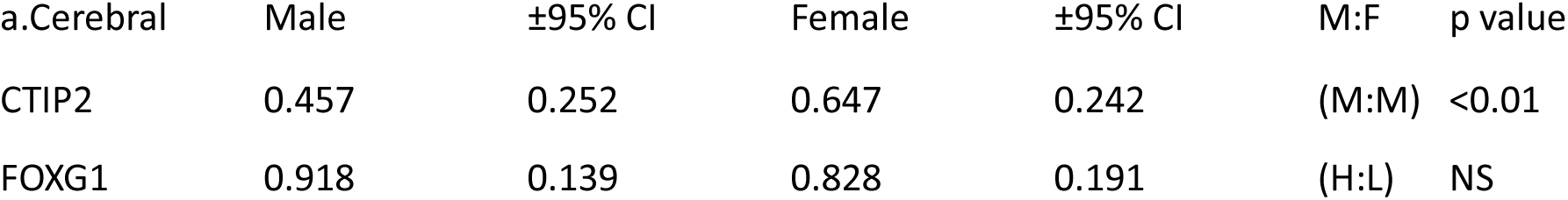
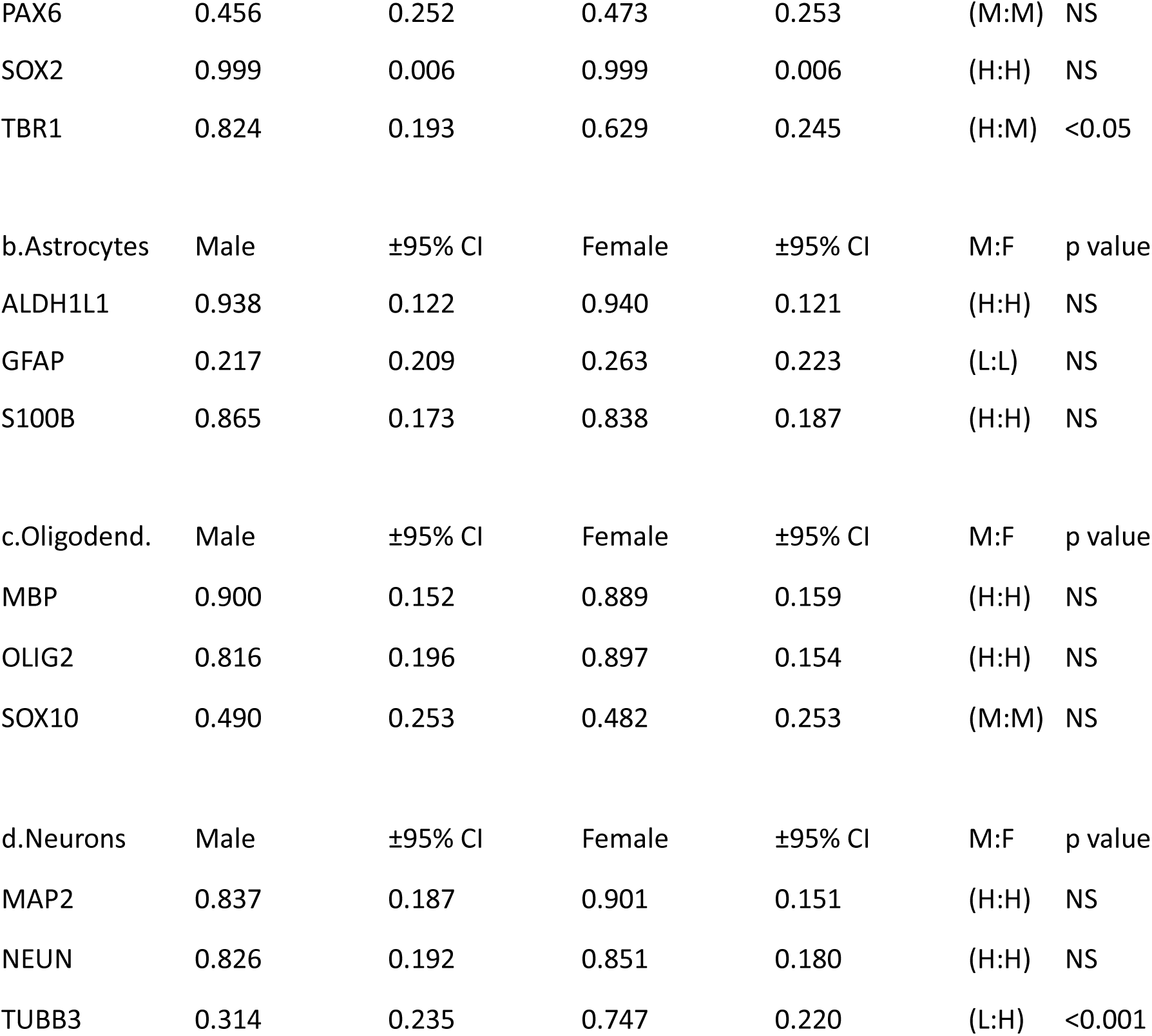
A comparison of male and female makers of a physiological CNS.

In summary, among the brain cell markers analyzed, CTIP2, TBR1, and TUBB3 showed significant gender differences, with CTIP2 and TBR1 being more expressed in females and males respectively, and TUBB3 showing a much higher expression in females. All other markers showed similar levels of expression between genders.

### 3. Brainstem

#### 3.1. Overview of the aiHumanoid Brainstem Simulation

In addition to the cerebrum, we also evaluated the brainstem component of the aiHumanoid CNS simulation. Here, we compared the expression levels of markers specific to serotonergic, noradrenergic, and cholinergic neurons in both male and female simulations.

#### 3.2. Serotonergic Neuron Marker (N=3) Expression in the aiHumanoid Simulation

The serotonergic neuron markers analyzed in this study included SERT (Serotonin Transporter), TPH2 (Tryptophan Hydroxylase 2, and VMAT2 (Vesicular Monoamine Transporter 2). While TPH2 and VMAT2 had similar expression levels in males and females, SERT expression levels were higher in females (p<0.001). These results are summarized in Table 3a.

#### 3.3. Noradrenergic Neuron (N=3) Marker Expression in the aiHumanoid Simulation

We assessed the expression of noradrenergic neuron markers, including DBH (Dopamine Beta-Hydroxylase), NET (Norepinephrine Transporter), and TH (Tyrosine Hydroxylase). TH expression levels were somewhat higher in females (p<0.05) while DBH and NET demonstrated similar expression in both genders (in Table 3b).

#### 3.4. Cholinergic Neuron Marker (N=3) Expression in the aiHumanoid Simulation

The cholinergic neuron markers evaluated included CHAT (Choline Acetyltransferase), CHT1 (Choline Transporter 1), and VAChT (Vesicular Acetylcholine Transporter). All three markers were expressed with no significant gender differences (in Table 3c).

**Table 3:**
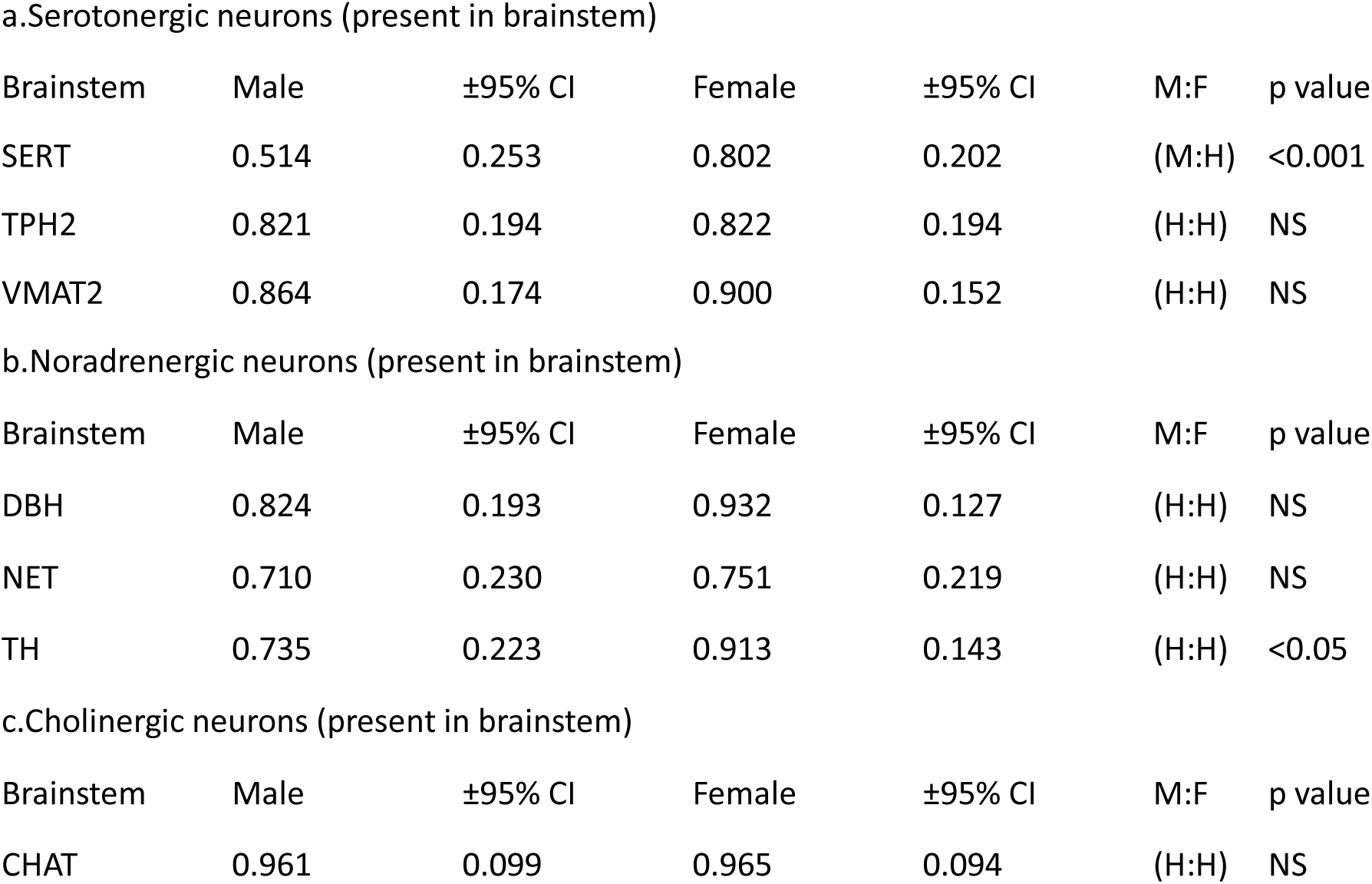
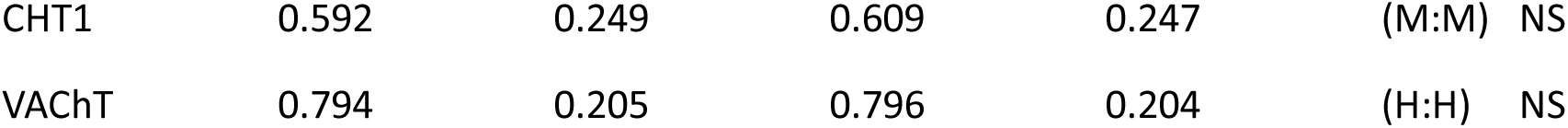
A comparison of male and female makers of a physiological Brainstem.

#### 3.5. Evaluation of the aiHumanoid Brainstem Simulation

The results demonstrate consistent expression of serotonergic, noradrenergic, and cholinergic neuron markers across both male and female brainstem simulations. Significant gender differences were only identified in SERT and TH markers.

### 4. Cerebellum

#### 4.1. Overview of the aiHumanoid Cerebellar Simulation Validation

In addition to cerebrum, and brainstem, we also explored the cerebellar component of the aiHumanoid CNS simulation. We compared the expression levels of markers specific to Purkinje cells, granule cells, Bergmann glia, molecular layer interneurons, GABAergic interneurons, and unipolar brush cells in both male and female simulations to evaluate the performance of the simulated cerebellum.

#### 4.2. Purkinje Cell Marker (N=3) Expression in the aiHumanoid Simulation

The Purkinje cell markers analyzed in this study included CALB1 (Calbindin 1), PCP2 (Purkinje Cell Protein 2), and RPL7 (Ribosomal Protein L7). All markers were expressed, and no gender differences were identified. These data are summarized in Table 4a.

#### 4.3. Granule Cell Marker (N=3) Expression in the aiHumanoid Simulation

We assessed the expression of granule cell markers, including NEUROD1 (Neurogenic Differentiation 1), PAX6 (Paired Box 6), and PROX1 (Prospero Homeobox 1). NEUROD1 and PAX6 were similarly expressed while PROX1 expression was higher in males (p<0.01) (in Table 4b).

#### 4.4. Bergmann Glia Marker (N=3) Expression in the aiHumanoid Simulation

The Bergmann glia markers evaluated included BLBP (Brain Lipid Binding Protein), GFAP (Glial Fibrillary Acidic Protein), and S100B (S100 Calcium Binding Protein B). All markers were expressed, and no gender differences were identified (in Table 4c).

#### 4.5. Molecular Layer Interneuron Marker (N=2) Expression in the aiHumanoid Simulation

The molecular layer interneuron markers analyzed included CALB2 (Calbindin 2) and PVALB (Parvalbumin). CALB2 expression levels were higher in females (p<0.001) but PVALB expression was similar in both genders (in Table 4d).

#### 4.6. GABAergic Interneuron Marker (N=2) Expression in the aiHumanoid Simulation

The GABAergic interneuron markers analyzed included GAD65 (Glutamate Decarboxylase 65) and GAD67 (Glutamate Decarboxylase 67). Both markers were expressed, and no gender differences were identified (in Table 4e)

#### 4.7. Unipolar Brush Cell Marker (N=2) Expression in the aiHumanoid Simulation

The unipolar brush cell markers evaluated included MET (MET Proto-Oncogene, Receptor Tyrosine Kinase) and PLCB4 (Phospholipase C Beta 4). MET expression was similar in both gender while PLCB4 was higher in females (p<0.001) (in Table 4f).

**Table 4:** A comparison of male and female markers of a physiological Cerebellum.

a.Purkinje cells
| Cerebellum | Male | ±95% CI | Female | ±95% CI | M:F | p value |
| --- | --- | --- | --- | --- | --- | --- |
| CALB1 | 0.784 | 0.208 | 0.819 | 0.195 | (H:H) | NS |
| PCP2 | 0.805 | 0.201 | 0.797 | 0.204 | (H:H) | NS |
| RPL7 | 0.805 | 0.201 | 0.797 | 0.204 | (H:H) | NS |

b.Granule cells
| Cerebellum | Male | ±95% CI | Female | ±95% CI | M:F | p value |
| --- | --- | --- | --- | --- | --- | --- |
| NEUROD1 | 0.668 | 0.238 | 0.729 | 0.225 | (M:H) | NS |
| PAX6 | 0.456 | 0.252 | 0.473 | 0.253 | (M:M) | NS |
| PROX1 | 0.626 | 0.245 | 0.848 | 0.182 | (M:H) | <0.01 |

c.Bergmann glia
| Cerebellum | Male | ±95% CI | Female | ±95% CI | M:F | p value |
| --- | --- | --- | --- | --- | --- | --- |
| BLBP | 0.844 | 0.184 | 0.928 | 0.131 | (H:H) | NS |
| GFAP | 0.217 | 0.209 | 0.263 | 0.223 | (L:L) | NS |
| S100B | 0.865 | 0.173 | 0.838 | 0.187 | (H:H) | NS |

d.Molecular layer interneurons
| Cerebellum | Male | ±95% CI | Female | ±95% CI | M:F | p value |
| --- | --- | --- | --- | --- | --- | --- |
| CALB2 | 0.313 | 0.235 | 0.536 | 0.252 | (L:M) | <0.001 |
| PVALB | 0.947 | 0.114 | 0.950 | 0.111 | (H:H) | NS |

e.GABAergic interneurons
| Cerebellum | Male | ±95% CI | Female | ±95% CI | M:F | p value |
| --- | --- | --- | --- | --- | --- | --- |
| GAD65 | 0.928 | 0.131 | 0.947 | 0.113 | (H:H) | NS |
| GAD67 | 0.440 | 0.251 | 0.421 | 0.250 | (M:M) | NS |

f.Unipolar brush cells
| Cerebellum | Male | ±95% CI | Female | ±95% CI | M:F | p value |
| --- | --- | --- | --- | --- | --- | --- |
| MET | 0.967 | 0.091 | 0.967 | 0.091 | (H:H) | NS |
| PLCB4 | 0.409 | 0.249 | 0.631 | 0.244 | (M:M) | <0.001 |

#### 4.8. Evaluation of the aiHumanoid Cerebellar Simulation

In summary, among the analyzed cerebellum cell markers, PROX1, CALB2, and PLCB4 showed significant gender differences, with all three markers showing higher expression in females. All other markers showed similar levels of expression between genders.

### 5. Gender specific organoids

#### 5.1 Overview of the Gender dependent aiHumanoid Simulations

Next, we explored the gender dependent simulations. We compared the expression levels of markers specific to mammary (breast) tissue, ovarian tissue, and testicular tissue, in both male and female simulations to evaluate the performance of the simulated organoids.

#### 5.2. Mammary Gland Marker (N=10) Expression in the aiHumanoid Simulation

For the mammary gland, the following markers were analyzed: CD24, CK14, KRT18, HER2, EPCAM, ER, GATA3, CD49f, CD29, and PR. Cluster of Differentiation 24 (CD24), Cytokeratin 14 (CK14), Human Epidermal growth factor Receptor 2 (HER2), Epithelial Cell Adhesion Molecule (EPCAM), GATA Binding Protein 3 (GATA3), Integrin Alpha 6 (CD49f), Integrin Beta 1 (CD29), Progesterone Receptor (PR), and Keratin 18 (KRT18) are significantly more expressed in females, except for Estrogen Receptor (ER) which shows no significant difference. GATA3 has a low level of expression in males and medium in females with a significant difference. These results are presented in Table 5a.

#### 5.3. Ovarian Marker (N=9) Expression in the aiHumanoid Simulation

For the ovarian tissue, the following markers were analyzed: AMH, CYP19A1, ER, FOXL2, FSHR, LHR, PAX8, PR, and WT1. Anti-Müllerian Hormone (AMH), Cytochrome P450 Family 19 Subfamily A Member 1 (CYP19A1), Forkhead Box L2 (FOXL2), and Progesterone Receptor (PR) are significantly more expressed in females. Estrogen Receptor (ER), Follicle Stimulating Hormone Receptor (FSHR), Luteinizing Hormone Receptor (LHR), Paired Box 8 (PAX8), and Wilms Tumor 1 (WT1) show no significant gender differences in expression levels (in Table 5b).

#### 5.4. Testicular Marker (N=10) Expression in the aiHumanoid Simulation

For the testicular tissue, the following markers were analyzed: AMH, DAZL, DMRT1, GATA4, PLZF, SOX9, SRY, SYCP3, Testosterone, and VASA. Anti-Müllerian Hormone (AMH), Doublesex and Mab-3 Related Transcription Factor 1 (DMRT1), Sex Determining Region Y (SRY), Testosterone (Test.), and VASA (also known as DDX4) have significantly higher expression in males. Deleted in Azoospermia-Like (DAZL), GATA Binding Protein 4 (GATA4), Promyelocytic Leukemia Zinc Finger (PLZF), and Synaptonemal Complex Protein 3 (SYCP3) show no significant gender difference, while SRY-Box Transcription Factor 9 (SOX9) shows a significant difference with higher expression in males (in Table 5c).

5.6. To summarize, the aiHumanoid simulation provides an initial validation of integrated organoid simulations with a common cardiovascular system. The data presented in this section demonstrates the expression levels and confidence intervals of specific markers for the various organoid simulations. These findings lay the groundwork for further exploration and refinement of this revolutionary simulation system. By analyzing the differences and similarities between male and female simulations, researchers can gain a better understanding of the complex interplay between these organoid systems and develop more sophisticated models in the future.

**Table 5:** A comparison of male and female makers of a physiological Gender specific organoids.

| a.Breast | Male | ±95% CI | Female | ±95% CI | M:F | p value |
| --- | --- | --- | --- | --- | --- | --- |
| CD24 | 0.599 | 0.248 | 0.749 | 0.219 | (M:H) | <0.05 |
| CK14 | 0.771 | 0.213 | 0.856 | 0.178 | (H:H) | NS |
| KRT18 | 0.537 | 0.252 | 0.869 | 0.171 | (M:H) | <0.001 |
| HER2 | 0.801 | 0.202 | 0.849 | 0.181 | (H:H) | NS |
| EPCAM | 0.335 | 0.239 | 0.755 | 0.218 | (M:H) | <0.001 |
| ER | 0.734 | 0.224 | 0.867 | 0.172 | (H:H) | NS |
| GATA3 | 0.148 | 0.180 | 0.664 | 0.239 | (L:M) | <0.001 |
| CD49f | 0.892 | 0.157 | 0.872 | 0.169 | (H:H) | NS |
| CD29 | 0.832 | 0.189 | 0.811 | 0.198 | (H:H) | NS |
| PR | 0.621 | 0.246 | 0.967 | 0.091 | (M:H) | <0.001 |
| b.Ovarian | Male | ±95% CI | Female | ±95% CI | M:F | p value |
| AMH | 0.965 | 0.093 | 0.555 | 0.252 | (H:M) | <0.001 |
| CYP19A1 | 0.538 | 0.252 | 0.996 | 0.030 | (M:H) | <0.001 |
| ER | 0.734 | 0.224 | 0.867 | 0.172 | (H:H) | NS |
| FOXL2 | 0.010 | 0.050 | 0.687 | 0.235 | (L:H) | <0.001 |
| FSHR | 0.145 | 0.178 | 0.164 | 0.187 | (L:L) | NS |
| LHR | 0.197 | 0.201 | 0.174 | 0.192 | (L:L) | NS |
| PAX8 | 0.903 | 0.150 | 0.934 | 0.126 | (H:H) | NS |
| PR | 0.621 | 0.246 | 0.967 | 0.091 | (M:H) | <0.001 |
| WT1 | 0.904 | 0.149 | 0.894 | 0.156 | (H:H) | NS |
| c.Testicular | Male | ±95% CI | Female | ±95% CI | M:F | p value |
| AMH | 0.965 | 0.093 | 0.555 | 0.252 | (H:M) | <0.001 |
| DAZL | 0.167 | 0.189 | 0.246 | 0.218 | (L:L) | NS |
| DMRT1 | 0.959 | 0.100 | 0.712 | 0.229 | (H:H) | <0.01 |
| GATA4 | 0.859 | 0.176 | 0.859 | 0.176 | (H:H) | NS |
| PLZF | 0.425 | 0.250 | 0.340 | 0.240 | (M:M) | NS |
| SOX9 | 0.962 | 0.096 | 0.749 | 0.220 | (H:H) | <0.05 |
| SRY | 0.871 | 0.170 | 0.305 | 0.232 | (H:L) | <0.001 |
| SYCP3 | 0.290 | 0.230 | 0.445 | 0.252 | (L:M) | <0.01 |
| Test. | 0.776 | 0.211 | 0.277 | 0.227 | (H:L) | <0.001 |
| VASA | 0.141 | 0.176 | 0.339 | 0.240 | (L:M) | <0.001 |

6.6 Overall, these data suggest that there are significant gender-specific differences in the expression levels of several key genes and proteins in breast, ovary, and testicular organoid simulations.

### 7. Endocrine

#### 7.1 Overview of the Endocrine aiHumanoid Simulations

Here we report on the data from the endocrine organoid simulations. We compared the expression levels of markers specific for the pancreas, thyroid, and adrenal glands in both male and females.

#### 7.2 Pancreas Marker(N=5) Expression in the aiHumanoid Simulation

For the pancreatic organoid, the following markers were analyzed: GCG (Glucagon), INS (Insulin), NKX6.1 (NK6 Homeobox 1), PDX1 (Pancreatic and Duodenal Homeobox 1), and SOX9 (SRY (Sex Determining Region Y)-Box 9). The expression levels of GCG, Insulin, and PDX1 were not statistically significant (NS) between the genders. However, the expression levels of NKX6.1 and SOX9 are higher in males than females as indicated by the statistically significant p-value of less than 0.05. These results are summarized in Table 7a.

#### 7.3 Thyroid Marker(N=5) Expression in the aiHumanoid Simulation

For the thyroid organoid, the following markers were analyzed: NIS (Sodium/Iodide Symporter), PAX8 (Paired Box 8), TG (Thyroglobulin), TPO (Thyroid Peroxidase), and TTF1 (Thyroid Transcription Factor 1). All five markers were expressed in both male and female simulations. Statistically significance gender differences in thyroid markers were not found. These results are summarized in Table 7b.

#### 7.4 Adrenal Marker(N=5) Expression in the aiHumanoid Simulation

For the adrenal gland organoid, the following markers were analyzed: CYP11B1 (Cytochrome P450 Family 11 Subfamily B Member 1), CYP11B2 (Cytochrome P450 Family 11 Subfamily B Member 2), CYP17A1 (Cytochrome P450 Family 17 Subfamily A Member 1), NR5A1 (Nuclear Receptor Subfamily 5 Group A Member 1), and STAR (Steroidogenic Acute Regulatory Protein). All five markers were expressed in both male and female simulations. Statistically significance gender differences in adrenal markers were not found. These results are also presented in Table 7c below.

**Table 7:** A comparison of male and female makers of a physiological Endocrine system.

| a.Pancreas | Male | ±95% CI | Female | ±95% CI | M:F | p value |
| --- | --- | --- | --- | --- | --- | --- |
| GCG | 0.251 | 0.219 | 0.253 | 0.220 | (L:L) | NS |
| INS | 0.915 | 0.141 | 0.888 | 0.160 | (H:H) | NS |
| NKX6.1 | 0.520 | 0.253 | 0.392 | 0.247 | (M:M) | <0.05 |
| PDX1 | 0.586 | 0.249 | 0.522 | 0.253 | (M:M) | NS |
| SOX9 | 0.962 | 0.096 | 0.749 | 0.220 | (H:H) | <0.05 |
| b.Thyroid | Male | ±95% CI | Female | ±95% CI | M:F | p value |
| NIS | 0.575 | 0.250 | 0.595 | 0.248 | (M:M) | NS |
| PAX8 | 0.903 | 0.150 | 0.934 | 0.126 | (H:H) | NS |
| TG | 0.805 | 0.201 | 0.816 | 0.196 | (H:H) | NS |
| TPO | 0.706 | 0.231 | 0.761 | 0.216 | (H:H) | NS |
| TTF1 | 0.882 | 0.163 | 0.862 | 0.175 | (H:H) | NS |
| c.Adrenal | Male | ±95% CI | Female | ±95% CI | M:F | p value |
| CYP11B1 | 0.432 | 0.251 | 0.402 | 0.248 | (M:M) | NS |
| CYP11B2 | 0.635 | 0.244 | 0.580 | 0.250 | (M:M) | NS |
| CYP17A1 | 0.832 | 0.189 | 0.886 | 0.161 | (H:H) | NS |
| NR5A1 | 0.961 | 0.098 | 0.913 | 0.142 | (H:H) | NS |
| STAR | 0.950 | 0.111 | 0.943 | 0.118 | (H:H) | NS |

7.5 In summary, while many of the endocrine markers show high expression levels in both sexes across the different glands, two markers in the pancreas (NKX6.1 and SOX9) showed a statistically significant difference (p<0.05) in expression between males and females. None of the other markers show significant gender-based differences in expression.

### 8. Internal Organoids

8.1 Gallbladder simulation results reveal varying marker (N=5) expression levels between male and female simulations. The CFTR (Cystic Fibrosis Transmembrane Conductance Regulator), KRT19 (Keratin 19), KRT7 (Keratin 7), MUC1 (Mucin 1, Cell Surface Associated), and MUC5AC (Mucin 5AC, Oligomeric Mucus/Gel-Forming). The data shows that gene expression levels for KRT19 and MUC5AC are significantly higher in female gallbladder organoids (p<0.001), while there are no significant differences for CFTR, KRT7, and MUC1 genes. These results are presented in Table 8a.

8.2 Intestinal simulation results exhibit some differences in marker (N=5) expression levels between male and female simulations. The markers evaluated were, the ASCL2 (Achaete-Scute Family BHLH Transcription Factor 2), CD44 (Cluster of Differentiation 44), EPHB2 (EPH Receptor B2), LGR5 (Leucine Rich Repeat Containing G Protein-Coupled Receptor 5), and OLFM4 (Olfactomedin 4). The results indicate significant differences in the expression of the CD44 and OLFM4 genes between male and female intestines. Both CD44 and OLFM4 were higher in the female organoids. The other markers (ASCL2, EPHB2, and LGR5) do not show significant gender-based differences in expression (in Table 8b).

8.3 Liver simulation results for N=4 markers; AFP (Alpha-Fetoprotein), CK19 (Cytokeratin 19), CK7 (Cytokeratin 7), and CYP3A4,5: Cytochrome P450 Family 3 Subfamily A Members 4,5). The data shows no significant differences in the expression of the AFP, CK19, CK7, and CYP3A4 genes between male and female livers. All expression levels are similar across genders (in Table 8c).

8.4 Kidney simulation results for N=5 markers reveal that the ECAD (E-Cadherin), LHX1 (LIM Homeobox 1), PAX2 (Paired Box 2), SIX2 (SIX Homeobox 2), and WT1 (Wilms Tumor 1) were all expressed. The data shows significant differences in the expression of ECAD, LHX1, PAX2, and SIX2 genes between male and female kidneys. Female organoid expression of these genes is consistently higher. However, there’s no significant difference in the expression of the WT1 gene between the genders (in Table 8d).

**Table 8:**
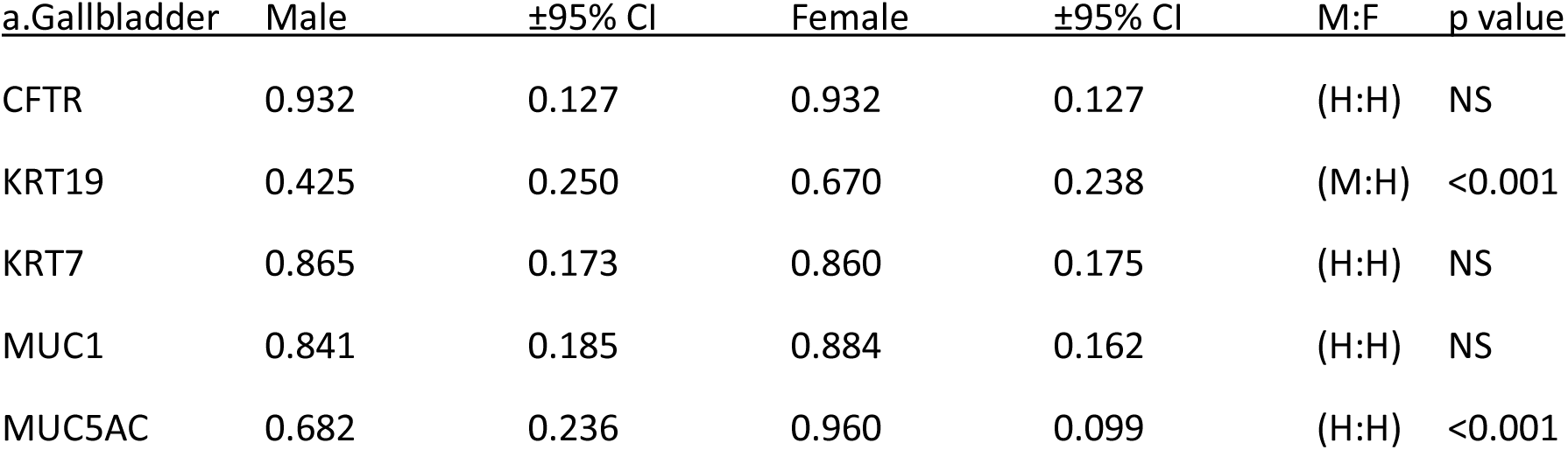
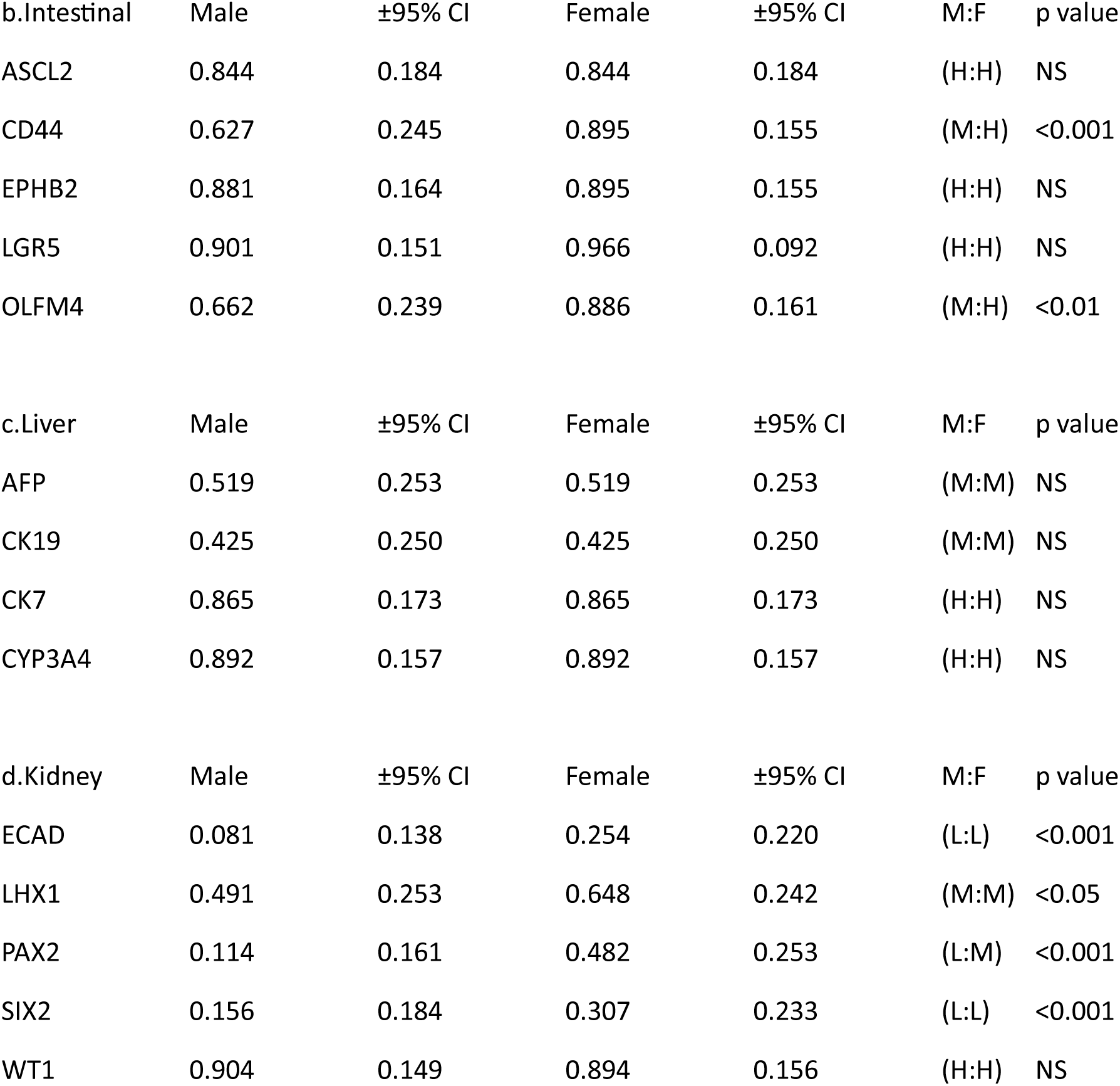
A comparison of male and female makers of a physiological Internal Organoids.

| <u>a.Gallbladder</u> | <u>Male</u> | <u>±95% CI</u> | <u>Female</u> | <u>±95% CI</u> | <u>M:F</u> | <u>p value</u> |
| --- | --- | --- | --- | --- | --- | --- |
| CFTR | 0.932 | 0.127 | 0.932 | 0.127 | (H:H) | NS |
| KRT19 | 0.425 | 0.250 | 0.670 | 0.238 | (M:H) | <0.001 |
| KRT7 | 0.865 | 0.173 | 0.860 | 0.175 | (H:H) | NS |
| MUC1 | 0.841 | 0.185 | 0.884 | 0.162 | (H:H) | NS |
| MUC5AC | 0.682 | 0.236 | 0.960 | 0.099 | (H:H) | <0.001 |

| b.Intestinal | Male | ±95% CI | Female | ±95% CI | M:F | p value |
| --- | --- | --- | --- | --- | --- | --- |
| ASCL2 | 0.844 | 0.184 | 0.844 | 0.184 | (H:H) | NS |
| CD44 | 0.627 | 0.245 | 0.895 | 0.155 | (M:H) | <0.001 |
| EPHB2 | 0.881 | 0.164 | 0.895 | 0.155 | (H:H) | NS |
| LGR5 | 0.901 | 0.151 | 0.966 | 0.092 | (H:H) | NS |
| OLFM4 | 0.662 | 0.239 | 0.886 | 0.161 | (M:H) | <0.01 |
| c.Liver | Male | ±95% CI | Female | ±95% CI | M:F | p value |
| AFP | 0.519 | 0.253 | 0.519 | 0.253 | (M:M) | NS |
| CK19 | 0.425 | 0.250 | 0.425 | 0.250 | (M:M) | NS |
| CK7 | 0.865 | 0.173 | 0.865 | 0.173 | (H:H) | NS |
| CYP3A4 | 0.892 | 0.157 | 0.892 | 0.157 | (H:H) | NS |
| d.Kidney | Male | ±95% CI | Female | ±95% CI | M:F | p value |
| ECAD | 0.081 | 0.138 | 0.254 | 0.220 | (L:L) | <0.001 |
| LHX1 | 0.491 | 0.253 | 0.648 | 0.242 | (M:M) | <0.05 |
| PAX2 | 0.114 | 0.161 | 0.482 | 0.253 | (L:M) | <0.001 |
| SIX2 | 0.156 | 0.184 | 0.307 | 0.233 | (L:L) | <0.001 |
| WT1 | 0.904 | 0.149 | 0.894 | 0.156 | (H:H) | NS |

8.5. In summary, some significant gender-based differences in gene expression were found in the gallbladder, intestines, and kidneys, but not the liver. The specific genes with significant differences varied by organoid.

### 9. Other

9.1 Lung simulation results reveal differences in marker (N=5)expression levels between male and female simulations. For instance, FOXJ1 (Forkhead Box J1), MUC5AC (Mucin 5AC, Oligomeric Mucus/Gel-Forming), NKX2.1 (NK2 Homeobox 1), P63 (Tumor Protein P63), and SOX9 (SRY (Sex Determining Region Y)-Box 9). The data shows significant differences in the expression of the FOXJ1, MUC5AC, and SOX9 genes between male and female lungs. FOXJ1 and MUC5AC expression were higher in females while SOX9 was higher in males. However, the NKX2.1 and P63 genes do not show significant gender-based differences in expression. These results are presented in Table 9a.

9.2 Skin simulation results display minimal variations in marker (N=5) expression levels between male and female simulations. The IVL (Involucrin), KRT14 (Keratin 14), KRT5 (Keratin 5), Loricrin (Loricrin Cornified Envelope Precursor Protein), and P63 (Tumor Protein P63). The data shows no significant differences in the expression of the IVL, KRT14, KRT5, LOR, and P63 genes between male and female skin. All expression levels are similar across genders (in Table 9b).

9.3 GUT-Brain axis (Normal)

For the GUT-Brain axis, the following markers (N=5) were analyzed: BDNF (Brain Derived Neurotrophic Factor), Cortisol, CRP (C-Reactive Protein), Haptoglobin (Zonulin) and SCFA (Short chain fatty acids). All five markers were expressed in both male and female simulations. No statistically significance gender differences in GUT-Brain markers were not found. These results are also presented in Table 9c below.

**Table 9.**
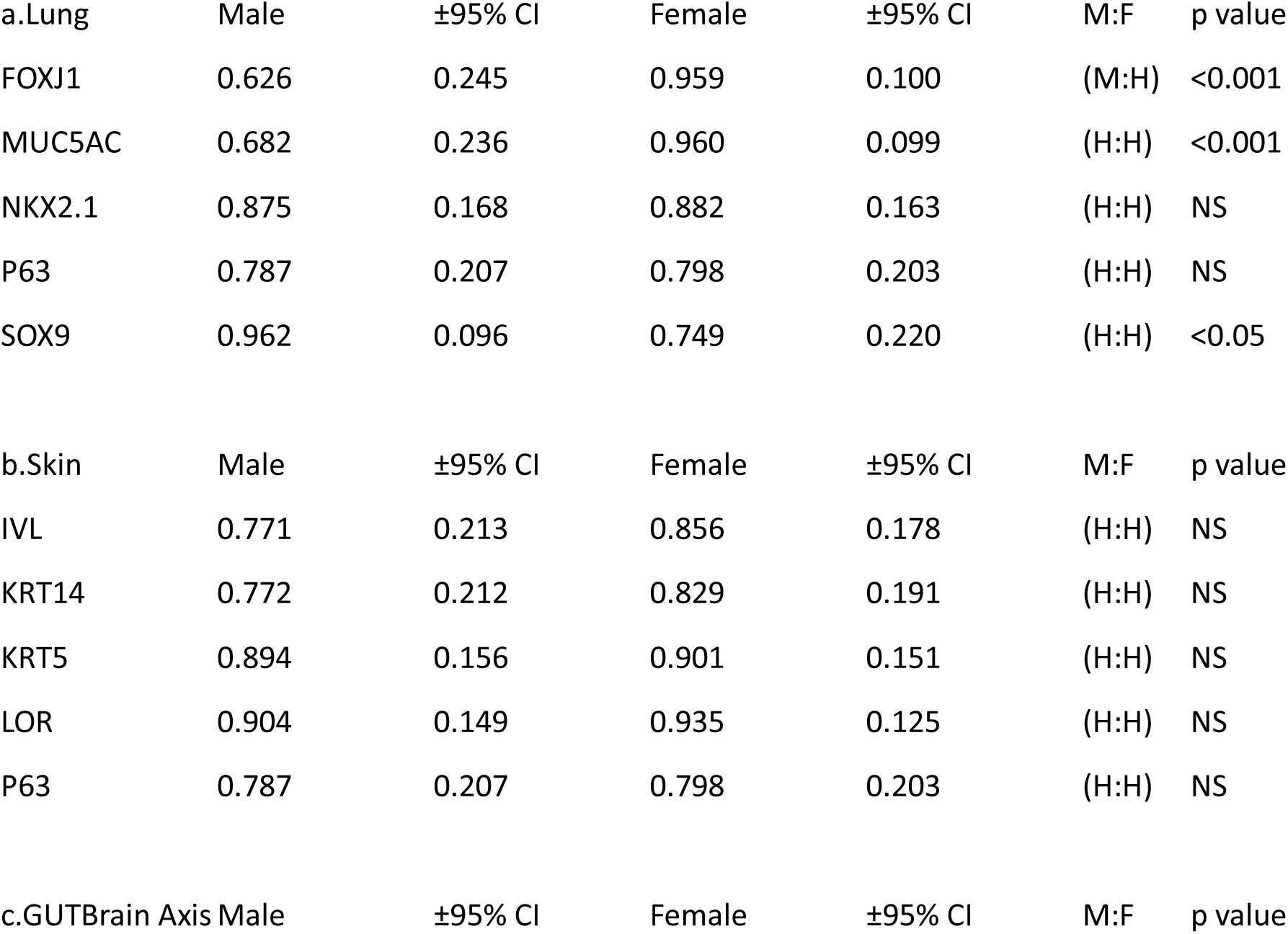
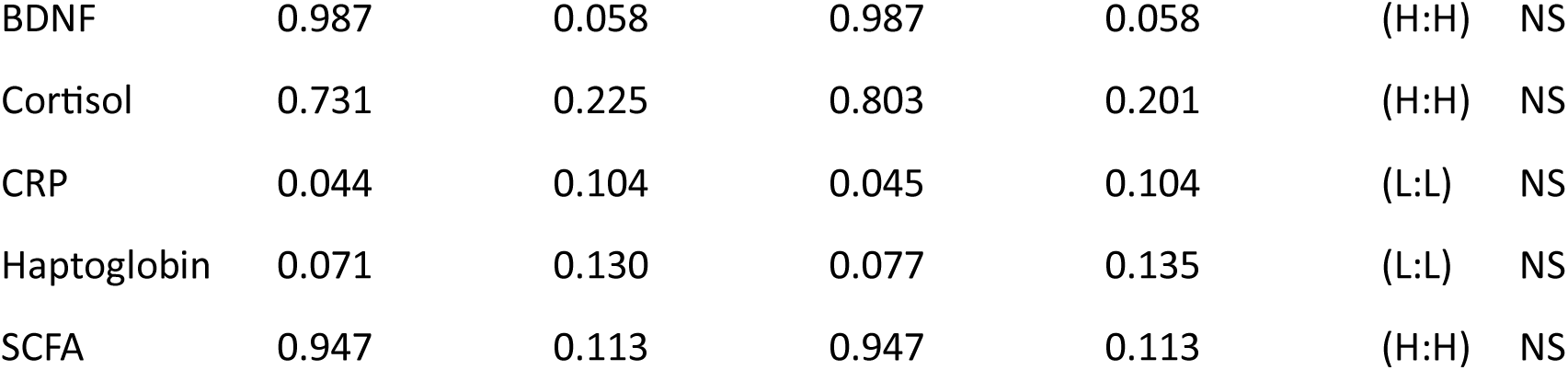

#### Overall Results Summary

The AI simulation of a multi-organoid humanoid has successfully demonstrated the potential to anticipate organoid interactions and predict their collective behavior. The model effectively incorporated both micro-level behavior of individual organoids and macro-level interactions within the system, thereby capturing a holistic view often missing in traditional in vitro models [28]. However, the accuracy of our model was found to be heavily dependent on the quality and extent of the input data, underscoring the importance of high-quality, comprehensive data in AI applications [29].

Despite demonstrating promising potential in simulating the general behavior of a multi-organoid system, the AI model faced challenges in predicting precise biological responses due to the inherent variability in biological systems (30). Moreover, the model’s interpretability in terms of underlying biological mechanisms needs further enhancement. Moving forward, the integration of AI with organoid technology, as evidenced by our study, could revolutionize a wide array of biomedical applications, notwithstanding the limitations and ethical considerations that need to be addressed [31].

## Discussion-Part 1

This research project was divided into two parts. In Part 1 we evaluated multiple markers for the uninfected aiHumanoid simulations to confirm that all recognized markers were present, representing all included organoid types. In Part 1, we also looked for evidence for gender differences in expression levels of specific markers for 18 organoid simulations. In Part 2 of the project, we explored the impact of gram-negative (like Pseudomonas aeruginosa) septicemia on the aiHumanoid simulation. Once again potential gender differences between males and females were considered in the analysis.

Cardiovascular System (CVS): The cardiac markers ACTC1, MYH6, MYH7, NKX2.5, and TNNT and the vascular markers aSMA, CD31, NG2, VE-Cadherin [32], and VWF [33] were all expressed in the CVS organoid simulations.

Our analysis of the cardiac markers revealed a significant difference in the expression of MYH7, with the male organoids showing a somewhat higher expression (p<0.05). The MYH7 gene encodes the beta-myosin heavy chain, a protein integral to cardiac muscle contraction [34]. The observed difference aligns with previous studies indicating gender-related differences in cardiac muscle physiology and response to stress [35]. The remainder of the cardiac markers such as ACTC1, MYH6, NKX2.5, and TNNT2 did not display significant gender-based differences, which is consistent with the suggestion that these genes play fundamental roles in cardiac physiology irrespective of gender [36].

On the other hand, the vascular marker, α-Smooth Muscle Actin (aSMA) showed a significantly higher expression in female organoids, hinting at potential gender-specific variations in vascular physiology. aSMA is a marker of vascular smooth muscle cells, which are key players in vascular function and disease [37]. The literature to date is quite limited regarding gender differences in aSMA expression, and this finding could lead to further investigation in this area.

Other vascular markers, including CD31, NG2, VE-Cadherin, and VWF, did not exhibit significant differences between male and female organoids. These markers are critical for various aspects of vascular function, including endothelial cell-cell junctions (CD31 and VE-Cadherin), pericyte coverage (NG2), and coagulation (VWF), and their similar expression in both sexes suggests these functions are preserved regardless of gender [38–41].

Overall, the expression of cardiac and vascular markers is consistent with the development of a CVS simulation which is central to engineering an integrated aiHumanoid. The possible gender differences in MYH7 and aSMA requires further investigation.

### The Central Nervous System

#### Cerebrum

The first component of the CNS to be evaluated was the physiologically normal cerebrum. The cerebral markers included those markers associated with cerebral cells, astrocytes, oligodendrocytes, and neurons. Once again, we uncovered intriguing differences between the male and female organoids, which could contribute to the growing body of evidence concerning gender-specific variations in brain physiology and function.

In the cerebral marker group, significant differences were found in the expression of CTIP2 and TBR1 between male and female organoids. CTIP2, a crucial gene involved in neuronal differentiation and axonal projection, showed a medium level of expression with significantly higher levels in female organoids [42]. TBR1, a gene necessary for glutamatergic neuronal differentiation, was highly expressed in both sexes but showed significantly higher expression in male organoids [43]. These findings could implicate gender-specific differences in neuronal differentiation and function, which warrant further investigation.

The Astrocyte markers ALDH1L1, GFAP, and S100B did not show significant gender-based differences, suggesting similar astrocytic functions in male and female brains [44]. Similarly, oligodendrocyte markers (MBP, OLIG2, SOX10) also demonstrated no significant differences, indicating comparable myelination processes across genders [45].

There were observed significant gender differences in the Neuron markers MAP2 and NEUN. However, the Neuronal marker TUBB3 did show a lower level of expression in male organoids and a high level in female organoids (p<0.001). TUBB3 participates in microtubule dynamics, which plays a key role in neuronal development and function [46]. This statistically significant gender-based difference could suggest potential variations in neuronal structures or functions between male and female brains.

In conclusion, the findings suggest potential gender-specific differences in cerebral physiology. These variations could provide insights into sex-based disparities in neurological and neuropsychiatric disorders and may guide the development of personalized therapeutic interventions. Future studies are warranted to elucidate the mechanisms underlying these differences and their potential clinical implications.

##### Brainstem

Next, we investigated the potential gender-specific differences in the normal physiologic markers of the Brainstem component of the CNS. These findings revealed significant differences in the expression of certain markers associated with serotonergic and noradrenergic neurons. No significant gender-based differences were observed in the markers of cholinergic brainstem neurons.

In the group of serotonergic neuron markers, SERT (Serotonin Transporter) exhibited a medium level of expression in males and a high level in females (p<0.001). SERT plays a crucial role in the reuptake of serotonin, a neurotransmitter implicated in mood regulation, sleep, and other functions [47]. The higher expression of SERT in females could suggest a gender-specific serotoninergic activity that might underpin differences in the prevalence and manifestation of certain neuropsychiatric disorders such as depression, which on a population basis seems to be more common in women [48].

For the noradrenergic neuron markers, TH (Tyrosine Hydroxylase), involved in the synthesis of norepinephrine, showed a significantly higher expression in females (p<0.05). This could indicate a gender-specific variation in the brainstem noradrenergic system, which is essential for the regulation of attention, arousal, and sleep-wake cycles [49].

Interestingly, no significant gender-based differences were found in the expression of cholinergic neuron markers (CHAT, CHT1, and VAChT), suggesting a comparable cholinergic activity in both sexes. This neurotransmitter system also plays a pivotal role in memory, attention, and arousal [50].

In conclusion, these data provide valuable insights into potential gender-specific differences in brainstem physiology. Better understanding these differences could shed light on gender disparities in various neurological and neuropsychiatric disorders and may inform the development of more effective gender-specific therapeutic interventions.

##### Cerebellum

The final component of the CNS examined potential gender-specific differences in normal physiologic markers for the Cerebellum. Here we also uncover findings suggesting some intriguing potential variations in cellular composition and function between males and females within specific cerebellar cell types.

The Purkinje cell markers, CALB1, PCP2, and RPL7 exhibited similar expression levels in both males and females, with no significant gender-based differences observed. Purkinje cells are the principal neurons of the cerebellar cortex, involved in motor coordination and cognitive functions [51]. The absence of significant gender-based differences in these markers suggests a comparable representation of Purkinje cells in the male and female cerebellum.

In the granule cell population, only PROX1 demonstrated potential gender-specific a gender difference. PROX1 exhibited a medium level of expression with significantly higher levels in females (p<0.01). Granule cells are the most abundant neuronal type in the cerebellum, participating in sensory integration and motor coordination [52]. The NeuroD1 and PAX6 markers were expressed at similar levels in both males and female simulations. The observed isolated difference in PROX1 expression is difficult to interpret but could suggest potential variations in granule cell development or function between genders.

Regarding Bergmann glia markers, BLBP, GFAP, and S100B showed similar expression levels in both males and females. Bergmann glia provide structural and metabolic support in the cerebellum and participate in neuronal migration and synaptic plasticity [53]. The absence of significant gender-based differences in these markers suggests a comparable representation of Bergmann glial cells in males and females.

In the molecular layer interneuron population, CALB2 displayed significantly lower expression in males compared to females (p<0.001), while PVALB showed no gender-based differences. Molecular layer interneurons play critical roles in modulating cerebellar circuitry and synaptic transmission [54]. The observed sex-based difference in CALB2 expression may imply potential disparities in molecular layer interneuron development or function between males and females, which could contribute to variations in cerebellar processing.

For the GABAergic interneuron markers, GAD65 and GAD67 exhibited no significant gender-based differences in their expression levels. GABAergic interneurons are key regulators of inhibitory neurotransmission in the cerebellum [55]. The absence of sex-based differences in GAD65 and GAD67 suggests a comparable representation of GABAergic interneurons in males and females.

Lastly, among the unipolar brush cell markers, MET showed similar expression levels between males and females, while PLCB4 exhibited significantly higher expression in females (p<0.001). Unipolar brush cells are specialized neurons involved in the modulation of excitatory input to Purkinje cells [56]. The observed sex-based difference in PLCB4 expression suggests potential sex-specific variations in the function or development of cerebellar unipolar brush cells.

In conclusion, these data provide insights into potential gender-specific differences in the cellular composition of the cerebellum. Understanding these differences may help to better understand gender-related disparities in cerebellar-related behaviors or disorders. Further research is warranted to elucidate the functional implications and underlying mechanisms of these observed variations.

##### Gender Specific Differences

To investigate potential differences in normal markers associated with gender-specific organoids, we focused on breast, ovarian, and testicular tissues. Our findings reveal significant variations in the expression of specific markers between males and females, highlighting the distinct molecular characteristics of these tissues in different sexes.

In the breast tissue, 5 of the 10 markers demonstrated significant gender-based differences. CD24 and KRT18 showed higher expression levels in females (p<0.05 and p<0.001 respectively), indicating their potential involvement in breast development and function [57,58]. GATA3 also exhibited significantly higher expression in females (p<0.001), suggesting its role in the regulation of mammary gland development and differentiation [59]. EPCAM (p<0.001) and PR (p<0.001) also showed higher expression levels in females, indicating their potential involvement in progesterone receptor signaling and hormone responsiveness [60,61]. Conversely, CK14, ER, CD29, CD48f and HER2 did not exhibit significant gender-based differences, suggesting their comparable expression in male and female organoids.

In the ovarian tissue, AMH and CYP19A1 displayed significant sex-based differences. AMH showed higher expression levels in males (p<0.001), consistent with its known role in testicular development and function [62]. CYP19A1 exhibited higher expression in females (p<0.001), consistent with its role in ovarian hormone biosynthesis [63]. FOXL2, a critical gene in ovarian development, also showed significantly higher expression in females (p<0.001), underscoring its essential role in female reproductive system development and maintenance [64]. PR expression was also increased in females as expected (p<0.001). The other markers, ER, FSHR, LHR, PAX8, and WT1 did not demonstrate significant gender-based differences, suggesting comparable expression levels in males and females.

In the testicular simulation, AMH, DMRT1, SOX9, SRY, and Testosterone exhibited significant gender-based differences. AMH displayed higher expression levels in males (p<0.001), consistent with its role in testicular development and the inhibition of female reproductive tract formation [62]. DMRT1 (p<0.01), SOX9 (p<0.05), and SRY (p<0.001) exhibited higher expression in males, highlighting their crucial roles in testicular differentiation and development [65,66,67]. SYCP3 (p<0.01) and VASA/DDX4 (p<0.001) showed significantly higher expression in females, indicating potential involvement in female reproductive functions [68,69]. As expected, Testosterone levels were markedly increased in the male (p<0.001) and much lower levels present in the female. PLZF and GATA4 did not exhibit significant gender-based differences, suggesting comparable expression levels between males and females.

In conclusion, our study provides valuable insights into the molecular characteristics of gender-specific organoids, highlighting significant sex-based differences in the expression of several specific markers. Understanding these differences may contribute to a better understanding of the development, function, and disorders associated with these tissues in males and females.

##### Endocrine Markers

All endocrine markers were expressed in all organoids. The analysis of these markers from the pancreas, thyroid, and adrenal gland simulations provided evidence for the differential expression of specific markers in males and females, highlighting the gender-based variations in endocrine tissue function and regulation.

In the pancreas, NKX6.1 exhibited significantly higher expression in males compared to females (p<0.05), suggesting a potential role in the regulation of pancreatic beta-cell development and insulin production [70]. SOX9 also displayed significantly higher expression in males, implying its involvement in the regulation of pancreatic exocrine cell development [71]. However, the expression levels of GCG, INS and PDX1 did not show significant gender-based differences, indicating comparable levels of glucagon and insulin production in males and females.

For the thyroid gland, the expression levels of NIS, PAX8, TG, TPO, and TTF1 did not demonstrate any significant gender-based differences. These markers play critical roles in thyroid hormone synthesis, transportation, and regulation [72,73]. The comparable expression levels between males and females suggest similar thyroid function and hormone production in both men and women.

For the adrenal glands, CYP11B1, CYP11B2, CYP17A1, NR5A1, and STAR expression did not exhibit significant gender-based differences. These markers participate in the synthesis and regulation of adrenal hormones, including glucocorticoids, mineralocorticoids, and sex steroids [74,75]. These results suggest comparable adrenal hormone production and regulation in both genders.

These findings highlight the importance of gender-specific markers in characterizing the endocrine organoids. While some markers demonstrated significant sex-based differences, suggesting potential gender-specific functional roles, others showed comparable expression levels between males and females, indicating common endocrine features shared by both sexes. Further studies are necessary to elucidate the underlying mechanisms and functional implications of these sex-based differences in endocrine tissues.

Understanding the sex-based variations in endocrine markers provides valuable insights into the development, function, and regulation of these tissues in males and females. It may contribute to a better understanding of gender-specific endocrine disorders, hormone-related diseases, and potential therapeutic interventions tailored to specific sexes.

##### Internal organoids

In the present study we also compared the expression of markers in internal organoids in males and females. The findings revealed distinct patterns of marker expression in different internal organs, indicating potential sex-based differences in organ function and regulation.

For example, in the gallbladder organoid, the expression levels of CFTR, KRT7, MUC1, and MUC5AC did not demonstrate significant gender-based differences. CFTR is a crucial transporter involved in chloride ion secretion, and its comparable expression levels between males and females suggest similar ion transport function in the gallbladder [76]. KRT7 and MUC1 are associated with epithelial cell integrity and mucin production, respectively, and their comparable expression levels indicate common structural and secretory features of the gallbladder in both genders [77,78]. However, KRT19 showed significantly higher expression in females (p<0.001), suggesting potential gender-specific roles in the regulation of gallbladder function [79]. MUC5AC also exhibited significantly higher expression in females (p<0.01), indicating a potential gender-specific contribution to the production of mucins in the gallbladder [80].

In the intestinal tissues, ASCL2, EPHB2, and LGR5 did not exhibit significant sex-based differences in their expression levels. These markers play essential roles in intestinal stem cell maintenance and proliferation [81,82,83]. The comparable expression levels suggest similar intestinal stem cell populations and regenerative capacity between males and females. However, CD44 (p<0.0010 and OLFM4 (p<0.01) both demonstrated significantly higher expression in females, suggesting potential gender-specific roles in intestinal cell adhesion and stem cell regulation [84,85].

In the liver, the expression levels of AFP, CK19, CK7, and CYP3A4 did not show significant gender-based differences. AFP is a marker of liver development, and its comparable expression levels suggest similar hepatic development between males and females [86]. CK19 and CK7 participate in maintaining the structural integrity of hepatocytes, and their comparable expression levels suggest similar hepatocyte function in both sexes [87]. CYP3A4 is a critical enzyme involved in drug metabolism in the liver, and its comparable expression levels suggest similar drug metabolism capacity between males and females [88].

In the kidney, ECAD, LHX1, PAX2, SIX2, and WT1 demonstrated distinct expression patterns in males and females. ECAD showed significantly lower expression in males (p<0.001), indicating potential gender-based differences in cell adhesion and epithelial integrity in the kidney [89]. LHX1 (p<0.05) and PAX2 (p<0.001) exhibited significantly higher expression in females, suggesting potential gender-specific roles in kidney development and differentiation [90,91]. SIX2 showed significantly lower expression in males (p<0.01), indicating potential gender-based differences in renal progenitor cell populations and renal development [92]. WT1 expression was not significantly different, suggesting comparable roles in kidney development and function between males and females [93].

These findings provide insights into the potential gender-based differences in marker expression within internal organs. While some markers showed comparable expression levels between males and females, suggesting similar organ function, others exhibited gender-based differences, indicating potential gender-specific roles in organ development, function, and regulation. Further research efforts are warranted to unravel the complex underlying mechanisms and functional implications of these gender-based differences in internal organ markers.

##### Other Organoids

This final section provides a comprehensive analysis of the findings presented in Table 9, comparing normal physiologic markers of various other organoids in males and females. Theses results highlight potential gender-specific differences in gene expression within the lungs, skin, and complex gut-brain axis.

In the lung organoid, the expression levels of FOXJ1 (p<0.001) and MUC5AC (p<0.001) were significantly higher in females compared to males, suggesting potential gender-based variations in lung function and mucin production [94,95]. However, the expression of NKX2.1 and P63 did not show significant gender differences, suggesting that these genes may play similar roles in both males and females in lung development and homeostasis [96]. SOX9 expression was higher in males (p<0.05) than females, suggesting a role for this transcription factor in gender-specific lung development or function. However, any gender differences in SOX9 lung expression remains to be proven [97].

Moving to the skin, the analysis revealed comparable expression levels of IVL, KRT14, KRT5, and LOR between males and females, suggesting that these genes may have similar functions in maintaining skin integrity and barrier function in both genders [98,99]. Similarly, P63 expression was not significantly different between males and females, indicating its crucial role in epidermal development and maintenance in both sexes [100].

Finally, the normal Gut-Brain Axis markers examined in this study included BDNF, cortisol, CRP, haptoglobin, and SCFA were expressed in both genders [101–106]. Interestingly, no significant gender-specific differences were observed in the expression levels of these markers. These results suggest that the gut-brain axis, which encompass neurotrophic factors, stress-related hormones, inflammatory markers, and microbial metabolites, may function similarly in both males and females.

Overall, this comparative analysis provides valuable insights into the potential gender-specific differences in gene expression in the lung organoids, skin organoids, and the gut-brain axis. Understanding these differences can contribute to a deeper understanding of organ development, function, and potential susceptibility to gender-specific diseases. Further research is needed to elucidate the underlying mechanisms and functional implications of these observed variations in gene expression. Additionally, considering the complex interplay between genetics, hormonal influences, and environmental factors is crucial in deciphering the full extent of gender-related differences in organ physiology and health. It is clear from the cumulative results that gender specific differences in many organoid systems is lacking.

While the Discussion so far has focused on gender differences in organoid specific expression, we should note the marked similarities when we compare 127 makers in 18 different organoids. Only 37 or 29.1% of the markers demonstrated variable gender-based expression. In other words, about 70% of the markers examined were comparably expressed in the male and female aiHumanoid. In addition, 12 of the 37 gender-specific markers were found in the gender-specific organoids (Breast, Ovary and Testicle) as expected.

### Part 2 – Gram-negative Sepsis

Gram negative sepsis is a life-threatening infection caused by gram-negative bacteria, like Pseudomonas aeruginosa, invading the bloodstream. It can lead to multiorgan failure, shock, and death if not treated promptly and effectively. Gram negative sepsis continues to be a major challenge for clinicians and researchers because it is associated with high mortality rates and increasing antibiotic resistance. Recent advances in the diagnosis, management, and prevention of gram-negative sepsis have been reviewed by [107–110].

In Part 2 of this study, we evaluated the impact of gram-negative sepsis on 11 different organoids of the aiHumanoid simulations. Those systems are CVS, Nervous system, Respiratory System, Renal system, Hepatic system, Hematologic system, Gastrointestinal system, Integumentary/MSK system, Immune system, Endocrine system, and Other. The large number of sepsis markers discussed below were used to assess and compare the genotypic and phenotypic response of male and female organoid simulations.

## Results-Part 2

10.1. The impact of gram-negative sepsis on the aiHumanoid Cardiovascular System (CVS) The CVS sepsis markers (N=8) evaluated included Troponin, ANF (Atrial Natriuretic Factor), BNP (B-type Natriuretic Peptide), Endothelin-1, Lactate, Arterial BP, Heart Rate and Contractility.

The column numbers represent the mean values of these parameters in the different groups. The 95% Confidence Interval (CI) provides an estimation of the uncertainty around these mean values. The p-value indicates the statistical significance of the difference between the WT and sepsis groups. The additional column (WT:S) provides the qualitative comparison of the mean values between the WT and sepsis-affected groups. This column provides a snapshot of how each parameter is affected in sepsis relative to the healthy state, thereby offering a quick visual aid for understanding the directional trend of the sepsis-induced changes.

a. Male Data: Troponin, Endothelin-1, Lactate, Arterial BP, and Heart Rate show a significant difference (p<0.001) between the WT and sepsis groups. Contractility shows a less robust but statistically significant difference (p<0.05). ANF and BNP, however, do not show a significant difference (NS) between the groups.

b. Female Data: Like the male data, Troponin, Endothelin-1, Lactate, Arterial BP, and Heart Rate show a significant difference (p<0.001) between the WT and sepsis groups. Contractility does not show a significant difference (NS, p=0.052), though the p-value is close to the conventional 0.05 threshold. Again, ANF and BNP do not show a significant difference (NS) between the groups. These results are summarized in Table 10a and 10b.

**Table 10:**
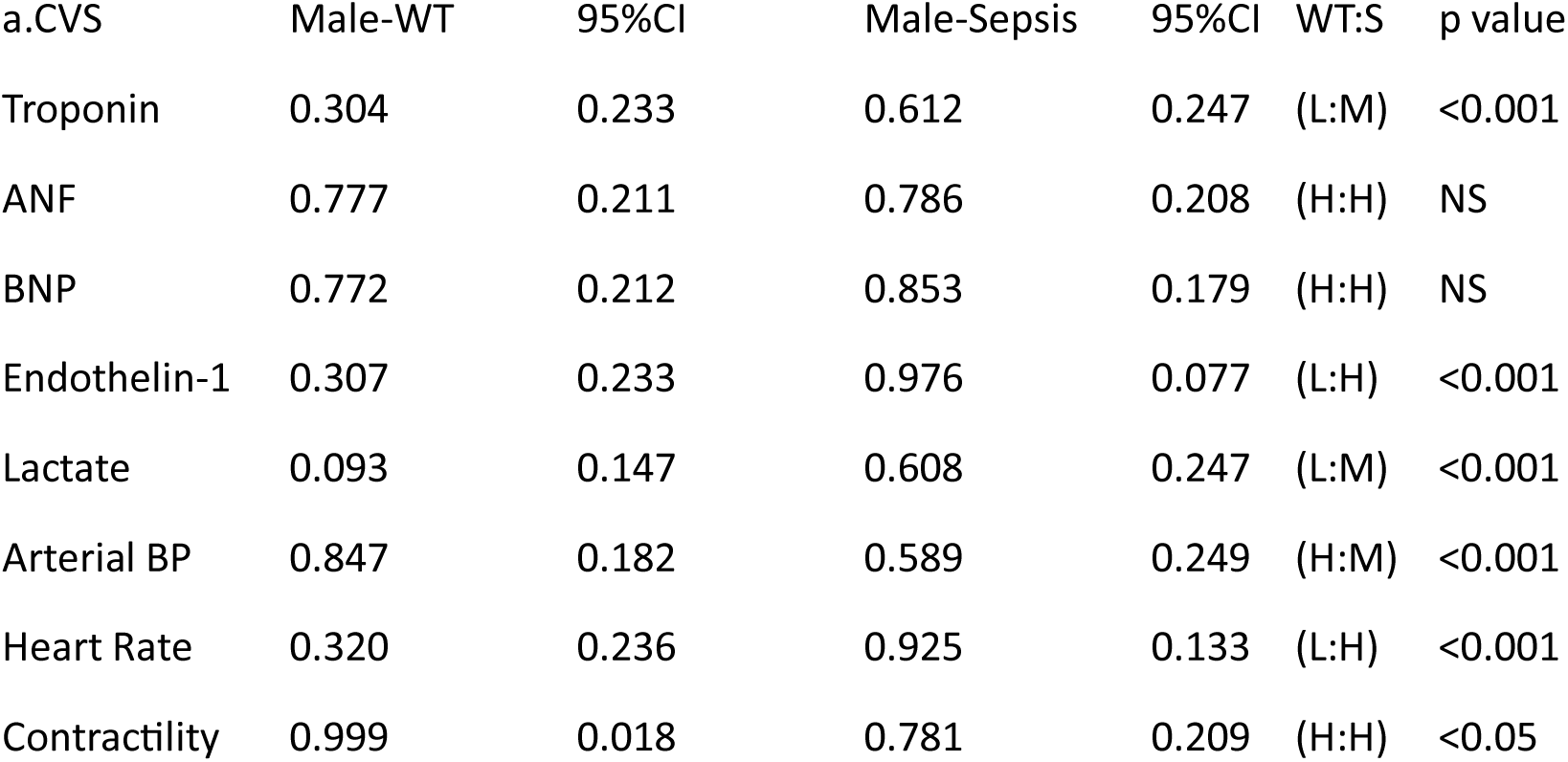
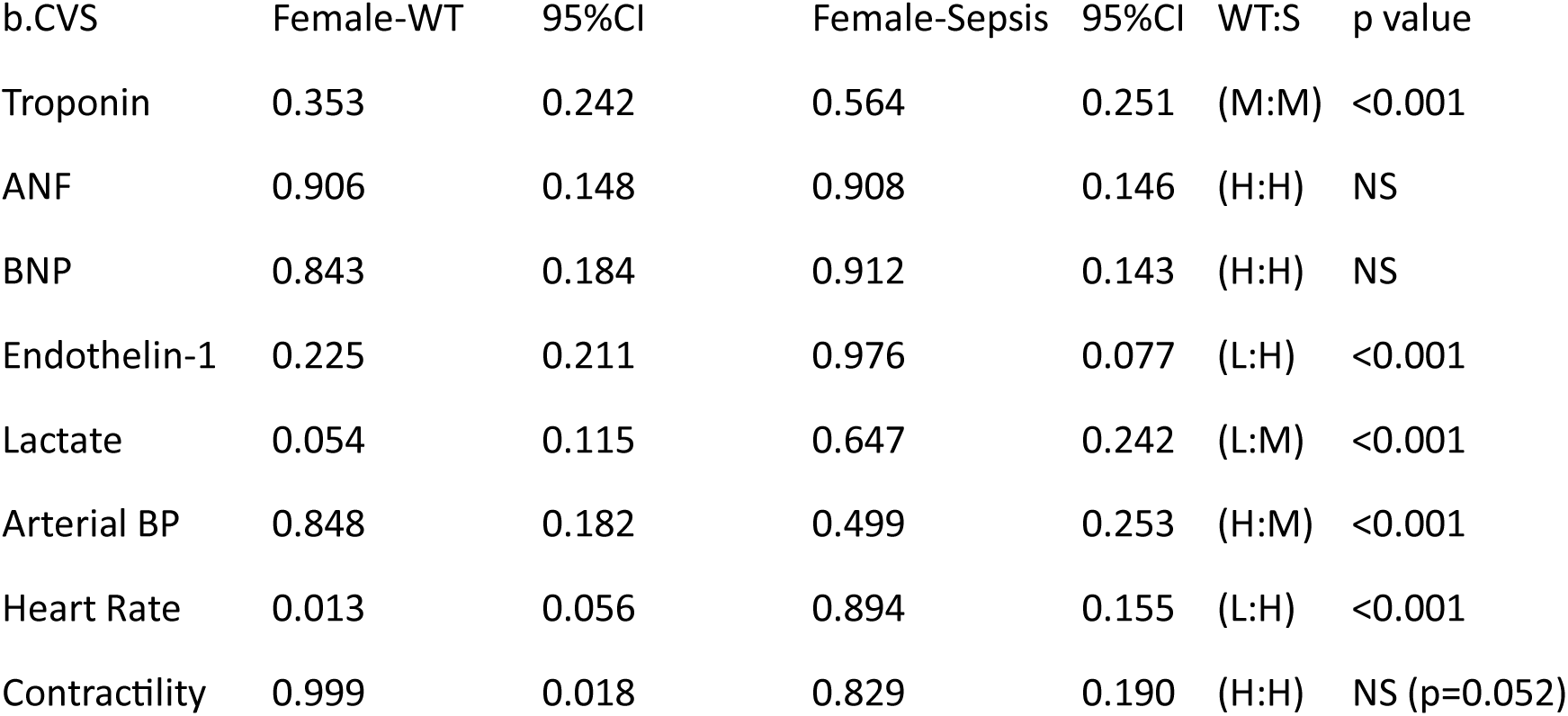
The impact of gram-negative sepsis on the aiHumanoid Cardiovascular System (CVS)

10.2. In summary, most cardiovascular parameters appear to be significantly altered in the sepsis condition compared to healthy individuals, both in males and females. However, the responses of ANF, BNP, and Contractility to sepsis appear to be more variable between genders.

11.1. The impact of gram-negative sepsis on the aiHumanoid Nervous System

The nervous system markers of gram-negative sepsis (N=3) included S100B (S100 Calcium Binding Protein B), NSE (Neuron-Specific Enolase), and GFAP (Glial Fibrillary Acidic Protein).

a. Male Data: NSE and GFAP show a significant difference (p<0.001) between the WT and sepsis groups. S100B, however, does not show a significant difference (NS) between the groups.

b. Female Data: Like the male data, NSE and GFAP show a significant difference (p<0.001) between the WT and sepsis groups. Again, S100B does not show a significant difference (NS) between the groups. These results are summarized in Table 11a and 11b.

**Table 11:** The impact of gram-negative sepsis on the aiHumanoid Nervous System.

| a.CNS | Male-WT | 95%CI | Male-Sepsis | 95%CI | WT:S | p value |
| --- | --- | --- | --- | --- | --- | --- |
| S100B | 0.901 | 0.151 | 0.933 | 0.127 | (H:H) | NS |
| NSE | 0.572 | 0.250 | 0.903 | 0.150 | (M:H) | <0.001 |
| GFAP | 0.172 | 0.191 | 0.958 | 0.101 | (L:H) | <0.001 |

| b.CNS | Female-WT | 95%CI | Female-Sepsis | 95%CI | WT:S | p value |
| --- | --- | --- | --- | --- | --- | --- |
| S100B | 0.862 | 0.175 | 0.911 | 0.144 | (H:H) | NS |
| NSE | 0.575 | 0.250 | 0.902 | 0.150 | (M:H) | <0.001 |
| GFAP | 0.292 | 0.230 | 0.958 | 0.101 | (L:H) | <0.001 |

11.2. In summary, most of the CNS parameters (NSE and GFAP) appear significantly altered in the sepsis condition compared to healthy individuals, both in males and females. However, the response of S100B to sepsis appears to be not significantly different between the two conditions, irrespective of gender.

12.1. The impact of gram-negative sepsis on the aiHumanoid Respiratory System

The respiratory markers of gram-negative sepsis (N=10) included SP-A (Surfactant proteins A), SP-D (Surfactant proteins D), CC16 (Clara cell secretory protein), sRAGE (Soluble Receptor for Advanced Glycation End-products), Alveolar Gas exchange, PaO2, PaCO2, Acute Lung injury, Respiratory failure, Respiratory Rate.

a. Male Data: SF-D, Gas Exchange, PaCO2, Acute Injury, Respiratory Failure, and Respiratory Rate show a significant difference between the WT and sepsis groups (p<0.05 or less). SF-A, CC16, sRAGE, and PaO2, however, do not show a significant difference (NS) between the groups.

b. Female Data: Gas Exchange, PaCO2, Lung Injury, Respiratory Failure, and Respiratory Rate show a significant difference between the WT and sepsis groups (p<0.01 or less). SF-A, SF-D, CC16, sRAGE, and PaO2, however, do not show a significant difference (NS) between the groups. These results are summarized in Table 12a and 12b.

**Table 12:** The impact of gram-negative sepsis on the aiHumanoid Respiratory System.

| a.Resp. System | Male-WT | 95%CI | Male-Sepsis | 95%CI | WT:S | p value |
| --- | --- | --- | --- | --- | --- | --- |
| SF-A | 0.770 | 0.213 | 0.649 | 0.242 | (H:M) | NS |
| SF-D | 0.765 | 0.215 | 0.607 | 0.247 | (H:M) | <0.05 |
| CC16 | 0.821 | 0.194 | 0.827 | 0.192 | (H:H) | NS |
| sRAGE | 0.782 | 0.209 | 0.908 | 0.146 | (H:H) | NS |
| Gas Exchange | 0.980 | 0.070 | 0.400 | 0.248 | (H:M) | <0.001 |
| PaO<sub>2</sub> | 0.887 | 0.160 | 0.846 | 0.183 | (H:H) | NS |
| PaCO<sub>2</sub> | 0.348 | 0.241 | 0.151 | 0.181 | (M:L) | <0.001 |
| Acute injury | 0.012 | 0.055 | 0.889 | 0.159 | (L:H) | <0.001 |
| Resp. failure | 0.101 | 0.152 | 0.967 | 0.091 | (L:H) | <0.001 |
| Resp. rate | 0.483 | 0.253 | 0.849 | 0.181 | (M:H) | <0.001 |

| b.Resp. System | Female-WT | 95%CI | Female-Sepsis | 95%CI | WT:S | p value |
| --- | --- | --- | --- | --- | --- | --- |
| SF-A | 0.744 | 0.221 | 0.730 | 0.225 | (H:H) | NS |
| SF-D | 0.685 | 0.235 | 0.619 | 0.246 | (H:M) | NS |
| CC16 | 0.786 | 0.208 | 0.803 | 0.202 | (H:H) | NS |
| sRAGE | 0.833 | 0.189 | 0.908 | 0.146 | (H:H) | NS |
| Gas exchange | 0.981 | 0.069 | 0.429 | 0.251 | (H:M) | <0.001 |
| PaO2 | 0.901 | 0.151 | 0.846 | 0.183 | (H:H) | NS |
| PaCO2 | 0.282 | 0.229 | 0.154 | 0.183 | (L:L) | <0.01 |
| Lung injury | 0.018 | 0.068 | 0.899 | 0.153 | (L:H) | <0.001 |
| Resp. failure | 0.124 | 0.167 | 0.967 | 0.091 | (L:H) | <0.001 |
| Resp. rate | 0.491 | 0.253 | 0.839 | 0.186 | (M:H) | <0.001 |

11.2. In summary, the sepsis condition appears to significantly alter several of the respiratory parameters in both males and females. However, some parameters, such as SF-A, SF-D, CC16, sRAGE, and PaO2, do not show a significant difference between healthy and sepsis-affected individuals, regardless of gender.

### The impact of gram-negative sepsis on the aiHumanoid Renal System

13.1. The renal markers of gram-negative sepsis (N=13) included NGAL (Neutrophil Gelatinase-Associated Lipocalin), KIM-1 (Kidney Injury Molecule-1), Cystatin C, Serum creatinine, BUN (Blood urea nitrogen, Urine Output, Acute kidney injury (AKI), GFR (Glomerular filtration rate), CKD/CRF (chronic kidney disease), Metabolic acidosis, Potassium, Sodium, and Proteinuria.

a. Male Data: All renal system parameters except Potassium and Sodium show a significant difference (p<0.001) between the WT and sepsis groups.

b. Female Data: Again, all parameters except Potassium and Sodium show a significant difference (p<0.01 or less) between the WT and sepsis groups. These results are summarized in Table 13a and 13b.

**Table 13:**
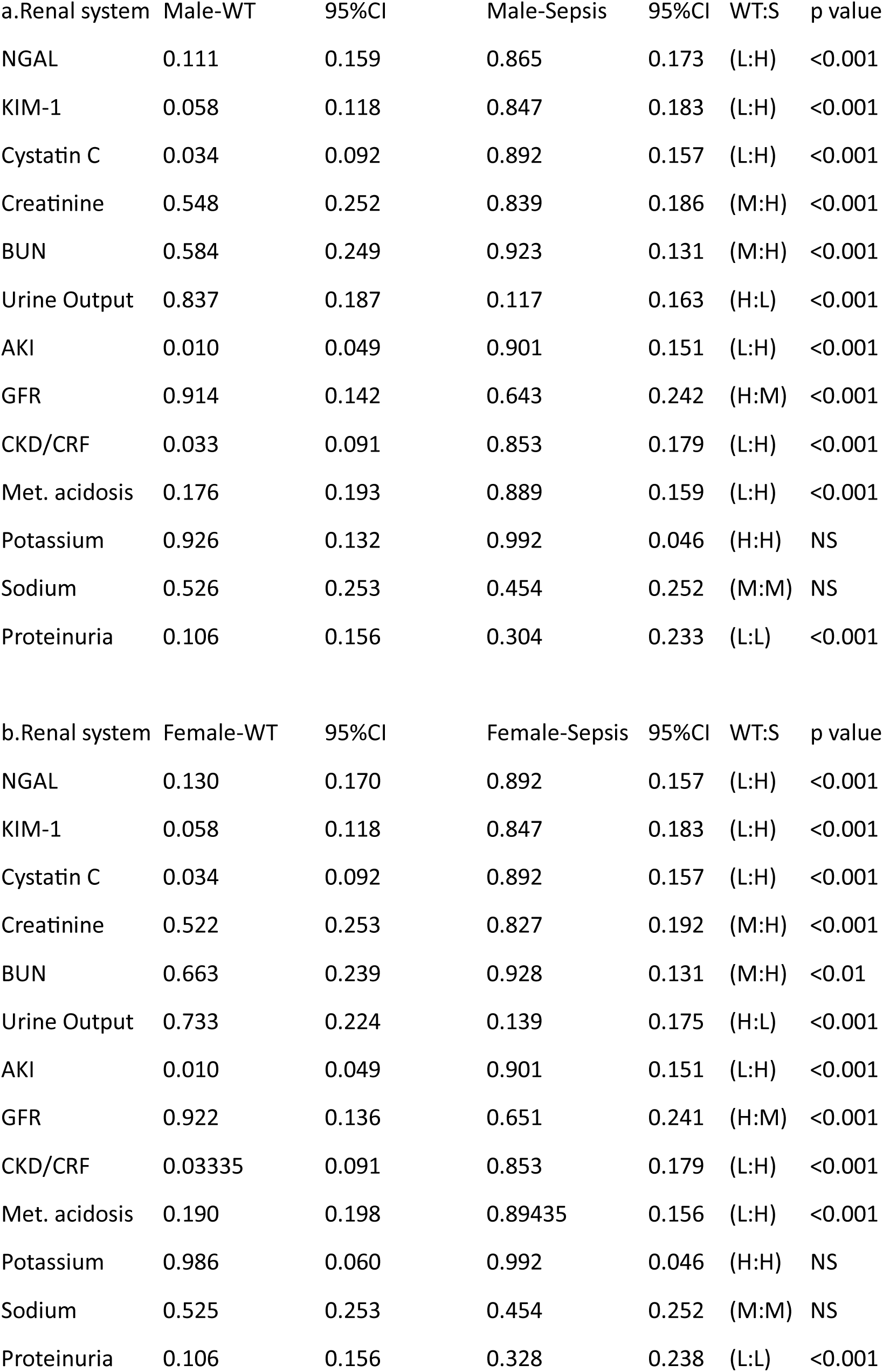
The impact of gram-negative sepsis on the aiHumanoid Renal System.

| a.Renal system | Male-WT | 95%CI | Male-Sepsis | 95%CI | WT:S | p value |
| --- | --- | --- | --- | --- | --- | --- |
| NGAL | 0.111 | 0.159 | 0.865 | 0.173 | (L:H) | <0.001 |
| KIM-1 | 0.058 | 0.118 | 0.847 | 0.183 | (L:H) | <0.001 |
| Cystatin C | 0.034 | 0.092 | 0.892 | 0.157 | (L:H) | <0.001 |
| Creatinine | 0.548 | 0.252 | 0.839 | 0.186 | (M:H) | <0.001 |
| BUN | 0.584 | 0.249 | 0.923 | 0.131 | (M:H) | <0.001 |
| Urine Output | 0.837 | 0.187 | 0.117 | 0.163 | (H:L) | <0.001 |
| AKI | 0.010 | 0.049 | 0.901 | 0.151 | (L:H) | <0.001 |
| GFR | 0.914 | 0.142 | 0.643 | 0.242 | (H:M) | <0.001 |
| CKD/CRF | 0.033 | 0.091 | 0.853 | 0.179 | (L:H) | <0.001 |
| Met. acidosis | 0.176 | 0.193 | 0.889 | 0.159 | (L:H) | <0.001 |
| Potassium | 0.926 | 0.132 | 0.992 | 0.046 | (H:H) | NS |
| Sodium | 0.526 | 0.253 | 0.454 | 0.252 | (M:M) | NS |
| Proteinuria | 0.106 | 0.156 | 0.304 | 0.233 | (L:L) | <0.001 |

| b.Renal system | Female-WT | 95%CI | Female-Sepsis | 95%CI | WT:S | p value |
| --- | --- | --- | --- | --- | --- | --- |
| NGAL | 0.130 | 0.170 | 0.892 | 0.157 | (L:H) | <0.001 |
| KIM-1 | 0.058 | 0.118 | 0.847 | 0.183 | (L:H) | <0.001 |
| Cystatin C | 0.034 | 0.092 | 0.892 | 0.157 | (L:H) | <0.001 |
| Creatinine | 0.522 | 0.253 | 0.827 | 0.192 | (M:H) | <0.001 |
| BUN | 0.663 | 0.239 | 0.928 | 0.131 | (M:H) | <0.01 |
| Urine Output | 0.733 | 0.224 | 0.139 | 0.175 | (H:L) | <0.001 |
| AKI | 0.010 | 0.049 | 0.901 | 0.151 | (L:H) | <0.001 |
| GFR | 0.922 | 0.136 | 0.651 | 0.241 | (H:M) | <0.001 |
| CKD/CRF | 0.03335 | 0.091 | 0.853 | 0.179 | (L:H) | <0.001 |
| Met. acidosis | 0.190 | 0.198 | 0.89435 | 0.156 | (L:H) | <0.001 |
| Potassium | 0.986 | 0.060 | 0.992 | 0.046 | (H:H) | NS |
| Sodium | 0.525 | 0.253 | 0.454 | 0.252 | (M:M) | NS |
| Proteinuria | 0.106 | 0.156 | 0.328 | 0.238 | (L:L) | <0.001 |

13.1. In summary, the sepsis condition appears to significantly alter most of the renal parameters in both males and females. However, some parameters such as Potassium and Sodium do not show a significant difference between healthy and sepsis-affected individuals, regardless of gender. These findings suggest that sepsis has a significant impact on renal function and may contribute to the development of conditions such as Acute Kidney Injury (AKI) and chronic kidney disease/Chronic Renal Failure (CKD/CRF).

### The impact of gram-negative sepsis on the aiHumanoid Hepatic System

14.1. The hepatic markers of gram-negative sepsis (N=8) included Albumin, ALT (Aspartate Aminotransferase), Alk. Phos. (Alkaline Phosphatase), AST (Alanine Aminotransferase), Bilirubin, Hepatocyte Infection, GGT (Gamma-Glutamyl Transferase), and Liver Damage/Cell Death.

a. Male Data: ALT, AST, Bilirubin, Liver Infection, and Liver Damage show a significant difference (p<0.05 or less) between the WT and sepsis groups. Albumin, Alk Phos, and GGT do not show a significant difference between the groups.

b. Female Data: ALT, AST, Bilirubin, Liver Infection, and Liver Damage show a significant difference (p<0.05 or less) between the WT and sepsis groups. Albumin, Alk Phos, and GGT do not show a significant difference between the groups. These results are summarized in Table 14a and 14b.

**Table 14:**
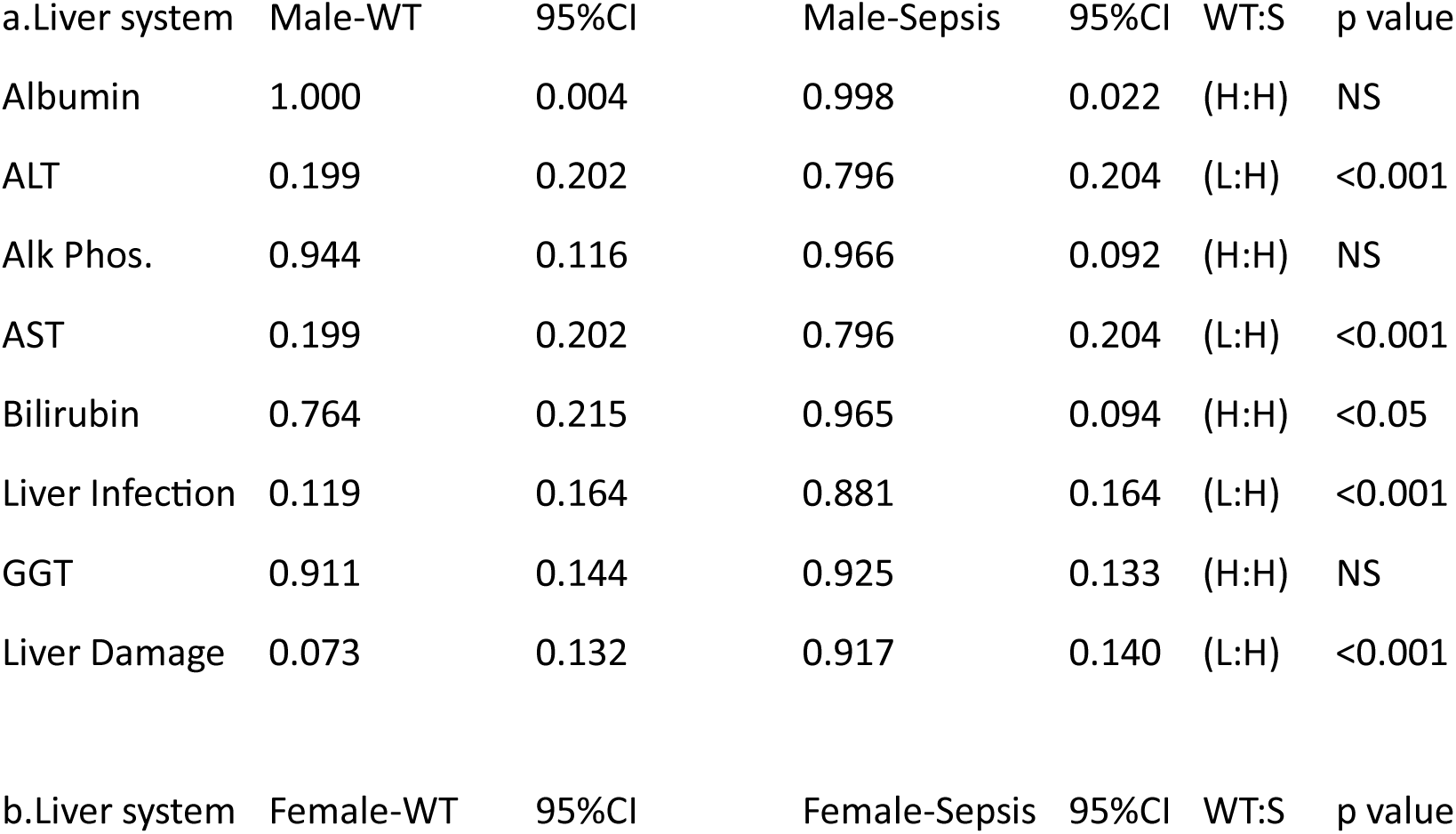
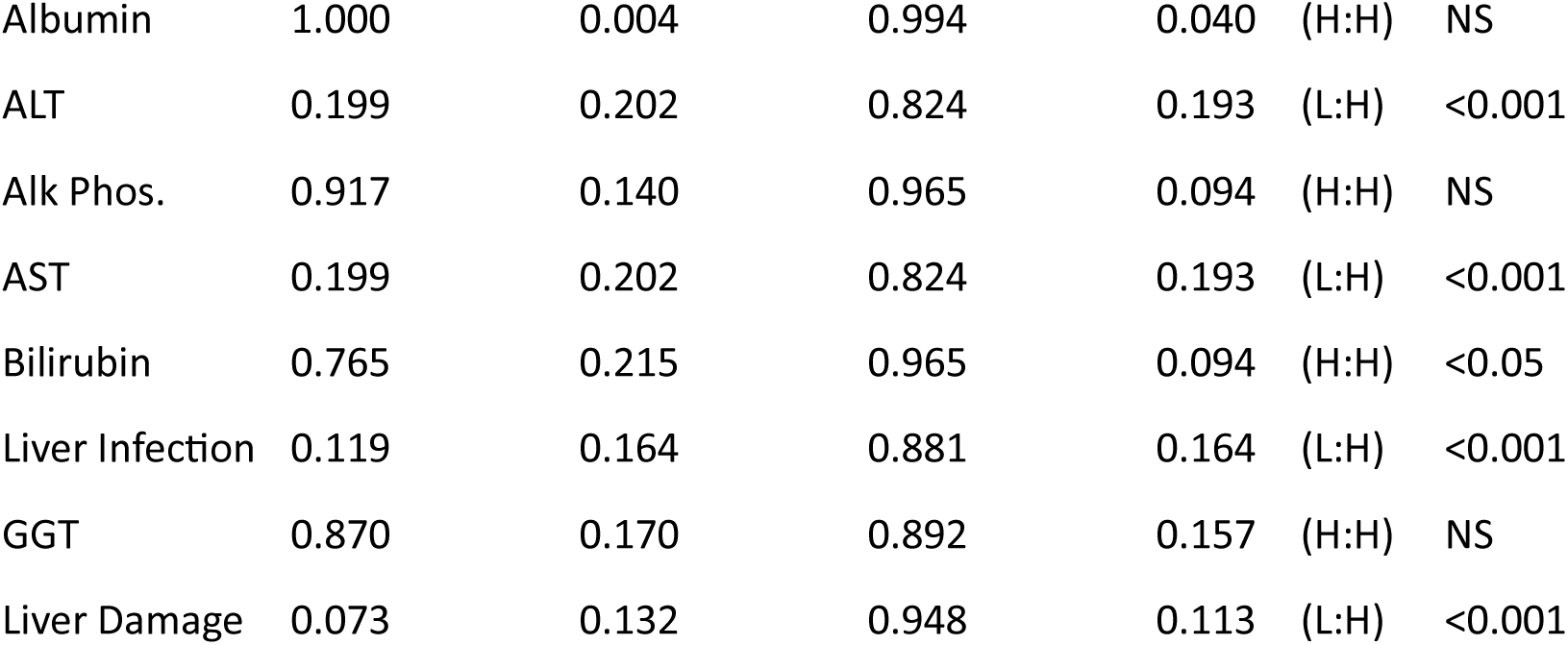
The impact of gram-negative sepsis on the aiHumanoid Hepatic System.

| a.Liver system | Male-WT | 95%CI | Male-Sepsis | 95%CI | WT:S | p value |
| --- | --- | --- | --- | --- | --- | --- |
| Albumin | 1.000 | 0.004 | 0.998 | 0.022 | (H:H) | NS |
| ALT | 0.199 | 0.202 | 0.796 | 0.204 | (L:H) | <0.001 |
| Alk Phos. | 0.944 | 0.116 | 0.966 | 0.092 | (H:H) | NS |
| AST | 0.199 | 0.202 | 0.796 | 0.204 | (L:H) | <0.001 |
| Bilirubin | 0.764 | 0.215 | 0.965 | 0.094 | (H:H) | <0.05 |
| Liver Infection | 0.119 | 0.164 | 0.881 | 0.164 | (L:H) | <0.001 |
| GGT | 0.911 | 0.144 | 0.925 | 0.133 | (H:H) | NS |
| Liver Damage | 0.073 | 0.132 | 0.917 | 0.140 | (L:H) | <0.001 |
| b.Liver system | Female-WT | 95%CI | Female-Sepsis | 95%CI | WT:S | p value |
| Albumin | 1.000 | 0.004 | 0.994 | 0.040 | (H:H) | NS |
| ALT | 0.199 | 0.202 | 0.824 | 0.193 | (L:H) | <0.001 |
| Alk Phos. | 0.917 | 0.140 | 0.965 | 0.094 | (H:H) | NS |
| AST | 0.199 | 0.202 | 0.824 | 0.193 | (L:H) | <0.001 |
| Bilirubin | 0.765 | 0.215 | 0.965 | 0.094 | (H:H) | <0.05 |
| Liver Infection | 0.119 | 0.164 | 0.881 | 0.164 | (L:H) | <0.001 |
| GGT | 0.870 | 0.170 | 0.892 | 0.157 | (H:H) | NS |
| Liver Damage | 0.073 | 0.132 | 0.948 | 0.113 | (L:H) | <0.001 |

14.1. In summary, the sepsis condition appears to significantly alter several of the liver parameters in both males and females. Specifically, ALT, AST, Bilirubin, and indications of Liver Infection and Liver Damage show significant increases in sepsis-affected individuals, suggesting a considerable impact of sepsis on liver function and integrity. However, parameters like Albumin, Alk Phos, and GGT do not show a significant difference between healthy and sepsis-affected individuals, regardless of gender.

### The impact of gram-negative sepsis on the aiHumanoid Hematologic System

15.1. The hematologic markers of gram-negative sepsis (N=8) included Anemia, Bleeding, D-dimers (unique fibrin degradation products), DIC (Disseminated Intravascular Coagulation), FDPs (Fibrin Degradation Products), Platelet count, P-selectin, and Thrombosis.

a. Male Data: All parameters show a significant difference (p<0.05 or less) between the WT and sepsis groups, except for D-dimers.

b. Female Data: All parameters show a significant difference (p<0.05 or less) between the WT and sepsis groups, except for D-dimers. These results are summarized in Table 15a and 15b.

**Table 15:** The impact of gram-negative sepsis on the aiHumanoid Hematologic System.

| a.Hem. System | Male-WT | 95%CI | Male-Sepsis | 95%CI | WT:S | p value |
| --- | --- | --- | --- | --- | --- | --- |
| Anemia | 0.034 | 0.092 | 0.607 | 0.247 | (L:M) | <0.001 |
| Bleeding | 0.335 | 0.239 | 0.799 | 0.203 | (M:H) | <0.001 |
| D-dimers | 0.683 | 0.236 | 0.770 | 0.213 | (H:H) | NS |
| DIC | 0.284 | 0.228 | 0.688 | 0.235 | (L:H) | <0.001 |
| FDPs | 0.302 | 0.232 | 0.766 | 0.214 | (L:H) | <0.001 |
| Platelet count | 0.846 | 0.183 | 0.579 | 0.250 | (H:M) | <0.001 |
| P-selectin | 0.103 | 0.154 | 0.840 | 0.186 | (L:H) | <0.001 |
| Thrombosis | 0.010 | 0.051 | 0.881 | 0.164 | (L:H) | <0.001 |

| b.Hem. system | Female-WT | 95%CI | Female-Sepsis | 95%CI | WT:S | p value |
| --- | --- | --- | --- | --- | --- | --- |
| Anemia | 0.034 | 0.092 | 0.675 | 0.237 | (L:H) | <0.001 |
| Bleeding | 0.235 | 0.215 | 0.774 | 0.212 | (L:H) | <0.001 |
| D-dimers | 0.649 | 0.242 | 0.754 | 0.218 | (M:H) | NS |
| DIC | 0.230 | 0.213 | 0.659 | 0.240 | (L:M) | <0.001 |
| FDPs | 0.228 | 0.212 | 0.752 | 0.219 | (L:H) | <0.001 |
| Platelet count | 0.926 | 0.133 | 0.727 | 0.225 | (H:H) | <0.05 |
| P-selectin | 0.259 | 0.222 | 0.841 | 0.185 | (L:H) | <0.001 |
| Thrombosis | 0.013 | 0.057 | 0.881 | 0.164 | (L:H) | <0.001 |

15.1. In summary, sepsis appears to significantly impact several hematological parameters in both males and females. Specifically, factors related to coagulation and platelet function, such as Bleeding, DIC, FDPs, Platelet count, P-selectin, and Thrombosis show significant changes in sepsis-affected individuals, suggesting a significant impact of sepsis on hemostasis and coagulation processes. Moreover, the prevalence of Anemia also shows a significant increase in sepsis-affected individuals. However, the levels of D-dimers do not show a significant difference between healthy and sepsis-affected individuals, regardless of gender.

### The impact of gram-negative sepsis on the aiHumanoid Gastrointestinal (GI) System

16.1. The GI markers of gram-negative sepsis (N=3) included I-FABP, D-lactate, and Citrulline.

a. Male Data: Two of the parameters (I-FABP, D-lactate) show a significant difference (p<0.001) between the WT and sepsis groups, but Citrulline shows no significant difference (NS).

b. Female Data: Like the male data, the parameters (I-FABP, D-lactate) show a significant difference (p<0.001) between the WT and sepsis groups, while Citrulline shows no significant difference (NS). These results are summarized in Table 16a and 16b.

**Table 16:** The impact of gram-negative sepsis on the aiHumanoid Gastrointestinal System.

| a.GI system | Male-WT | 95%CI | Male-Sepsis | 95%CI | WT:S | p value |
| --- | --- | --- | --- | --- | --- | --- |
| I-FABP | 0.257 | 0.221 | 0.913 | 0.143 | (L:H) | <0.001 |
| D-lactate | 0.408 | 0.249 | 0.874 | 0.168 | (M:H) | <0.001 |
| Citrulline | 0.928 | 0.131 | 0.799 | 0.208 | (H:H) | NS |
| b.GI system | Female-WT | 95%CI | Female-Sepsis | 95%CI | WT:S | p value |
| I-FABP | 0.367 | 0.244 | 0.884 | 0.162 | (M:H) | <0.001 |
| D-lactate | 0.414 | 0.249 | 0.875 | 0.168 | (M:H) | <0.001 |
| Citrulline | 0.967 | 0.090 | 0.799 | 0.203 | (H:H) | NS |

16.1. In summary, sepsis appears to significantly impact I-FABP and D-lactate levels in the GI system for both males and females. However, the levels of Citrulline in the GI system do not show a significant difference between healthy and sepsis-affected individuals, regardless of gender.

### The impact of gram-negative sepsis on the aiHumanoid Musculoskeletal System (MKS)

17.1. This data provides a comparative analysis of various parameters related to the musculoskeletal (MSK) system between healthy (wild-type, WT) and sepsis-affected subjects, divided by gender (Male and Female). The parameters under study (N=4) include E-selectin (Endothelial-leukocyte adhesion molecule 1), sICAM-1 (Soluble Intercellular Adhesion Molecule-1), MMP8 (Matrix Metallopeptidase 8), and MMP9 (Matrix Metallopeptidase 9)

a. Male Data: E-selectin and sICAM-1 show a significant increase (p<0.001) in the sepsis group compared to the WT group. MMP8 shows a statistically significant difference (p<0.05) between the WT and sepsis groups, being higher in the sepsis group. MMP9 does not show a significant difference (NS) between the two groups.

b. Female Data: Like the male data, E-selectin and sICAM-1 show a significant increase (p<0.01 and p<0.001, respectively) in the sepsis group compared to the WT group. MMP8 shows a statistically significant difference (p<0.05) between the WT and sepsis groups, like the male data. MMP9 does not show a significant difference (NS) between the two groups, consistent with the male data. These results are summarized in Table 17a and 17b.

**Table 17:**
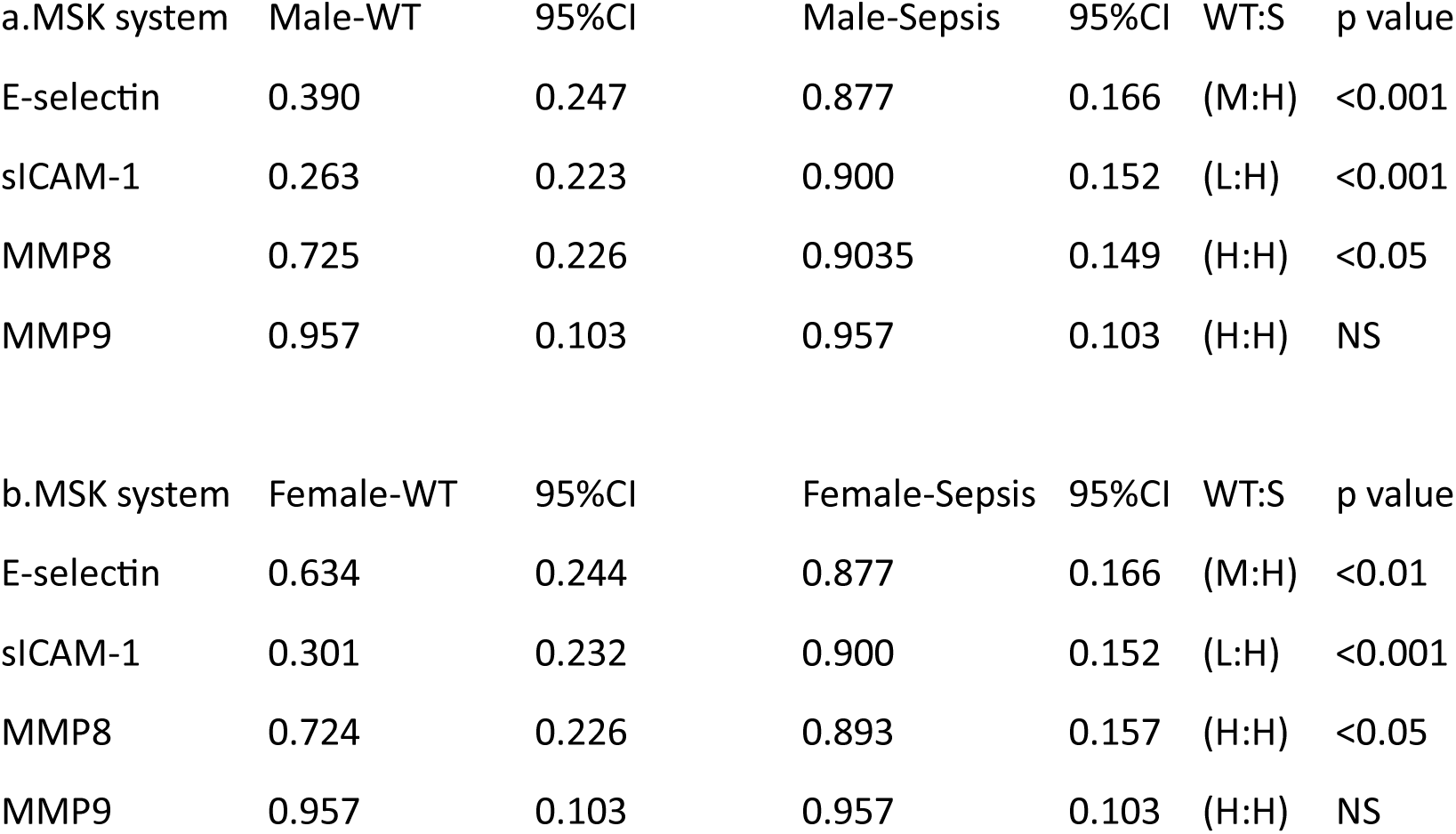
The impact of gram-negative sepsis on the aiHumanoid Musculoskeletal System.

17.1. In summary, sepsis appears to significantly impact E-selectin, sICAM-1, and MMP8 levels in the MSK system for both males and females, but MMP9 expression is not affected.

### The impact of gram-negative sepsis on the aiHumanoid Immune System

18.1. The Immune system markers of gram-negative sepsis (N=6) included IL-1b (Interleukin 1b), IL-6 (Interleukin 6), IL-8 (Interleukin 8), IL-10 (Interleukin 10), Chronic inflammation, and immunosuppression.

a. Male Data: All parameters listed (IL-1b, IL-6, IL-8, IL-10, Inflammation, Suppression) show a significant increase (p<0.001) in the sepsis group compared to the WT group.

b. Female Data: Like the male data, all parameters listed (IL-1b, IL-6, IL-8, IL-10, Inflammation, Suppression) show a significant increase (p<0.001) in the sepsis group compared to the WT group. These results are summarized in Table 18a and 18b.

**Table 18:** The impact of gram-negative sepsis on the aiHumanoid Immune Response.

| a.Immunity | Male-WT | 95%CI | Male-Sepsis | 95%CI | WT:S | p value |
| --- | --- | --- | --- | --- | --- | --- |
| IL-1b | 0.100 | 0.152 | 0.901 | 0.151 | (H:H) | <0.001 |
| IL-6 | 0.403 | 0.248 | 0.967 | 0.091 | (M:H) | <0.001 |
| IL-8 | 0.106 | 0.156 | 0.998 | 0.022 | (L:H) | <0.001 |
| IL-10 | 0.292 | 0.230 | 0.992 | 0.045 | (L:H) | <0.001 |
| Inflammation | 0.034 | 0.091 | 0.967 | 0.091 | (L:H) | <0.001 |
| Suppression | 0.403 | 0.248 | 0.967 | 0.091 | (M:H) | <0.001 |
| b.Immunity | Female-WT | 95%CI | Female-Sepsis | 95%CI | WT:S | p value |
| IL-1b | 0.100 | 0.152 | 0.901 | 0.151 | (L:H) | <0.001 |
| IL-6 | 0.377 | 0.245 | 0.967 | 0.091 | (M:H) | <0.001 |
| IL-8 | 0.189 | 0.198 | 0.998 | 0.022 | (L:H) | <0.001 |
| IL-10 | 0.372 | 0.245 | 0.992 | 0.045 | (M:H) | <0.001 |
| Inflammation | 0.034 | 0.091 | 0.967 | 0.091 | (L:H) | <0.001 |
| Suppression | 0.377 | 0.245 | 0.967 | 0.091 | (M:H) | <0.001 |

18.1. In summary, these results indicate that sepsis significantly alters the immune response in both males and females. These changes include an increase in certain cytokines (IL-1b, IL-6, IL-8, IL-10), indicative of an inflammatory response, and an increase in markers of immune suppression.

### The impact of gram-negative sepsis on the aiHumanoid Endocrine System

19.1. The Endocrine system markers of gram-negative sepsis (N=5) included Cortisol, Insulin, Adiponectin, Leptin, and C-peptide.

a. Male Data: Cortisol levels significantly increased (p<0.05) in the sepsis group compared to the WT group. Insulin and Adiponectin did not show a significant difference between the groups. Leptin and C-peptide levels showed a significant decrease (p<0.001) in the sepsis group compared to the WT group.

b. Female Data: Cortisol and Insulin did not show a significant difference between the groups. Adiponectin showed a significant decrease (p<0.001) in the sepsis group compared to the WT group. Leptin and C-peptide levels also showed a significant decrease (p<0.001) in the sepsis group compared to the WT group. These results are summarized in Table 19a and 19b.

**Table 19.** The impact of gram-negative sepsis on the aiHumanoid Endocrine System.

| a.Endocrine | Male-WT | 95%CI | Male-Sepsis | 95%CI | WT:S | p value |
| --- | --- | --- | --- | --- | --- | --- |
| Cortisol | 0.824 | 0.193 | 1.000 | 0.001 | (H:H) | <0.05 |
| Insulin | 0.939 | 0.121 | 1.000 | 0.004 | (H:H) | NS |
| Adiponectin | 0.945 | 0.115 | 0.837 | 0.187 | (H:H) | NS |
| Leptin | 0.143 | 0.177 | 0.714 | 0.229 | (L:H) | <0.001 |
| C-peptide | 0.587 | 0.249 | 0.322 | 0.237 | (M:L) | <0.001 |
| b.Endocrine | Female-WT | 95%CI | Female-Sepsis | 95%CI | WT:S | p value |
| Cortisol | 0.848 | 0.182 | 1.000 | 0.000 | (H:H) | NS |
| Insulin | 0.950 | 0.111 | 1.000 | 0.004 | (H:H) | NS |
| Adiponectin | 0.945 | 0.115 | 0.648 | 0.242 | (H:M) | <0.001 |
| Leptin | 0.158 | 0.185 | 0.741 | 0.222 | (L:H) | <0.001 |
| C-peptide | 0.572 | 0.250 | 0.308 | 0.234 | (M:L) | <0.001 |

19.1. In summary, these results suggest that sepsis significantly alters the endocrine response in both males and females. Cortisol, a hormone associated with stress, showed an increase in males with sepsis, but not in females. Both sexes exhibited a decrease in Leptin and C-peptide levels in the sepsis group, which could be indicative of metabolic alterations associated with sepsis.

### The impact of gram-negative sepsis on Other important aiHumanoid Systems

20.0. The Other important system markers of gram-negative sepsis (N=6) included Presepsin, PCT (ProCalcitonin), CRP (C-reactive protein), MODS (Multiple organ dysfunction syndrome), Catabolic state, and the Normal GUT-Brain Axis.

a. Male Data: Presepsin did not show a significant difference between the groups. PCT, CRP, MODS, and Catabolic state showed a significant increase (p<0.001) in the sepsis group compared to the WT group. Gut-Brain N showed a significant decrease (p<0.001) in the sepsis group compared to the WT group.

b. Female Data: Presepsin did not show a significant difference between the groups. PCT, CRP, MODS, and Catabolic state showed a significant increase (p<0.001) in the sepsis group compared to the WT group. Gut-Brain N showed a significant decrease (p<0.001) in the sepsis group compared to the WT group. These results are summarized in Table 20a and 20b.

**Table 20.** The impact of gram-negative sepsis on the aiHumanoid Other Systems.

| a.Other | Male-WT | 95%CI | Male-Sepsis | 95%CI | WT:S | p value |
| --- | --- | --- | --- | --- | --- | --- |
| Presepsin | 0.656 | 0.240 | 0.784 | 0.208 | (M:H) | NS |
| PCT | 0.076 | 0.134 | 0.964 | 0.094 | (L:H) | <0.001 |
| CRP | 0.056 | 0.116 | 0.966 | 0.092 | (L:H) | <0.001 |
| MODS | 0.133 | 0.172 | 0.933 | 0.127 | (L:H) | <0.001 |
| Catabolic state | 0.172 | 0.191 | 0.976 | 0.077 | (L:H) | <0.001 |
| GUT-Brain N | 0.965 | 0.094 | 0.038 | 0.096 | (H:L) | <0.001 |
| b.Other | Female-WT | 95%CI | Female-Sepsis | 95%CI | WT:S | p value |
| Presepsin | 0.627 | 0.245 | 0.762 | 0.216 | (M:H) | NS |
| PCT | 0.077 | 0.135 | 0.964 | 0.09 | (L:H) | <0.001 |
| CRP | 0.056 | 0.116 | 0.966 | 0.092 | (L:H) | <0.001 |
| MODS | 0.133 | 0.172 | 0.933 | 0.126 | (L:H) | <0.001 |
| Catabolic state | 0.186 | 0.197 | 0.976 | 0.077 | (L:H) | <0.001 |
| GUT-Brain N | 0.965 | 0.094 | 0.038 | 0.096 | (H:L) | <0.001 |

20.2. In summary, these results suggest that sepsis significantly impacts various biological parameters not strictly categorized under the main physiological systems. For example, PCT, CRP, and MODS, which are indicators of severe infection and organ dysfunction, are significantly increased in both males and females with sepsis. In contrast, the Gut-Brain N axis, shows a significant decrease in both sexes with sepsis. The data underscore the systemic and multifaceted impact of sepsis on human physiology.

### 21.1 Overall summary of the effects of sepsis on aiHumanoid organ systems

This comprehensive study uses aiHumanoid simulations to evaluate the impact of gram-negative sepsis on multiple organ systems, including the cardiovascular, nervous, respiratory, renal, hepatic, hematologic, gastrointestinal, musculoskeletal, and immune systems. The goal was to compare genotypic and phenotypic responses to gram-negative sepsis in male and female organoid simulations.

Most of the CVS markers, including Troponin, Endothelin-1, Lactate, Arterial BP, and Heart Rate were significantly altered in sepsis, with variable responses observed for ANF, BNP, and Contractility between genders. For the nervous system, NSE and GFAP were significantly affected, while S100B did not show a significant difference.

Respiratory system markers including SF-D, Gas Exchange, PaCO2, Acute Injury, Respiratory Failure, and Respiratory Rate were significantly impacted, while others like SF-A, CC16, sRAGE, and PaO2 remained unchanged.

Renal function was markedly affected, with all parameters showing a significant difference between healthy and sepsis-affected individuals, except for Potassium and Sodium. This suggests sepsis may contribute to conditions such as Acute Kidney Injury and Chronic Kidney Disease.

The hepatic system also showed significant alterations, with increased ALT, AST, Bilirubin, Liver Infection, and Liver Damage, while Albumin, Alk Phos, and GGT remained unaffected.

Gram-negative sepsis significantly impacted several hematological parameters, particularly those related to coagulation and platelet function, and increased the prevalence of Anemia. However, D-dimer levels remained unchanged.

In the gastrointestinal system, I-FABP and D-lactate levels were significantly impacted, while Citrulline levels remained unchanged. The musculoskeletal system showed significant changes in E-selectin, sICAM-1, and MMP8 levels, but not in MMP9.

Finally, sepsis significantly affected the immune response in both males and females, with marked increases in several cytokines such as IL-1b, IL-6, IL-8, IL-10, and evidence of chronic inflammation and immunosuppression.

These results offer important insights into the systemic impact of gram-negative sepsis, and the potential differences in organ system responses between genders, highlighting the complexity and diversity of the disease process.

## Part 2 - Discussion

The aiHumanoid simulation of gram-negative sepsis investigated the impact of sepsis on various physiological systems in comparison to wild-type (WT) conditions. The simulation results are consistent with the current understanding of sepsis pathophysiology, demonstrating widespread disruption of normal physiological processes and increased levels of markers associated with inflammation, organ dysfunction, and tissue damage [111].

For the CVS simulations, the sepsis group exhibited elevated levels of troponin, endothelin-1, lactate, and heart rate, reflecting increased cardiac stress, vasoconstriction, and impaired tissue perfusion [112]. Importantly, contractility was significantly lower in the sepsis group, indicating potential systolic dysfunction, which is a common finding in sepsis-associated cardiac dysfunction [113].

The Respiratory system in the sepsis group demonstrated reduced levels of surfactant proteins and impaired alveolar gas exchange, consistent with the development of acute lung injury and acute respiratory distress syndrome (ARDS) observed in septic patients [114]. Furthermore, elevated levels of sRAGE and acute lung injury markers suggest increased alveolar epithelial damage and inflammation [115].

Renal dysfunction was evident in the sepsis group, as demonstrated by increased levels of acute kidney injury markers (NGAL, KIM-1, cystatin C, and serum creatinine), suggesting that sepsis-induced acute kidney injury (AKI) is a significant contributor to morbidity and mortality [116]. Decreased urine output and GFR further support the presence of renal dysfunction [117].

Hepatic dysfunction in the sepsis group was evident from the elevated levels of hepatocytic enzymes (ALT and AST), bilirubin, and markers of hepatocyte infection and liver damage, indicating that sepsis can cause hepatic injury and impair liver function [118].

In the hematologic system, the sepsis group showed increased levels of anemia, bleeding, D-dimers, DIC, and thrombosis markers, suggesting a higher risk of coagulopathy and vascular complications in sepsis [119]. The nervous system also exhibited elevated levels of S100B, NSE, and GFAP, indicative of potential neurological damage and inflammation [120].

Gastrointestinal dysfunction was indicated by increased levels of I-FABP and D-lactate in the sepsis group, suggesting sepsis-associated gut barrier dysfunction and impaired enterocyte function [121]. The integumentary system results showed increased levels of E-selectin, sICAM-1, and MMP8, reflecting endothelial activation and tissue remodeling [122,123].

In the immune system, the sepsis group exhibited elevated levels of pro-inflammatory cytokines, acute-phase proteins, and markers of chronic inflammation and immunosuppression, consistent with the hyperinflammatory and immunosuppressive states observed in septic patients [124]. Finally, the endocrine system results demonstrated increased cortisol, CRP, and leptin levels, suggesting stress response and metabolic dysregulation in sepsis [125].

In conclusion, our aiHumanoid simulation of gram-negative sepsis demonstrates numerous significant impacts on multiple physiological systems, including cardiovascular, respiratory, renal, hepatic, hematologic, nervous, gastrointestinal, integumentary, immune, and endocrine systems. The findings are consistent with previous literature that highlights the widespread effects of sepsis and the potential for multiple organ dysfunction syndrome (MODS) [111,125–128].

The elevated levels of pro-inflammatory cytokines and acute-phase proteins in the sepsis group support the well-established role of systemic inflammation in sepsis pathophysiology. The observed alterations in markers of organ function and tissue damage corroborate the concept that sepsis can lead to organ dysfunction, which is a key component of sepsis diagnosis and management [129,130,131].

The use of an aiHumanoid in this study provides a controlled environment to investigate the complex interactions and responses that occur during gram-negative sepsis. However, it is important to acknowledge that the simulation may not fully capture the intricacies of real-life septic patients. Future research should focus on validating these findings in experimental and clinical settings, as well as exploring potential therapeutic targets to mitigate the detrimental effects of sepsis on various physiological systems.

Moreover, timely diagnosis and appropriate management of sepsis remain critical to improving patient outcomes, as sepsis is a significant global health concern with high morbidity and mortality rates [132–134]. The implementation of guidelines, such as the Surviving Sepsis Campaign [135,136], has shown promise in reducing sepsis-associated mortality and should continue to be a focus in sepsis research and clinical practice.

Soon, advanced technologies including organoids, AI-driven simulations, and nanotechnology [137] promise to significantly improve our understanding of sepsis pathophysiology. By facilitating the development of targeted therapeutic strategies, these innovations will alleviate the devastating impact of sepsis and substantially improve patient outcomes.

### Limitations and Future Directions

Attempting to create an aiHumanoid from multiple organoid simulations is an ambitious undertaking that is heavily dependent on the availability of high-quality data. While the DeepNEU database continues to evolve, it currently covers about 32% of the human genome and much more data and development will be required. The goal is to achieve ∼99% coverage over the next 3-5 years. Assuming we aim for 99% coverage of ∼22,500 genes, this should produce a minimum of ∼506 million neurons in the aiHumanoid simulations. While still modest in the context of a human being, it may be more realistic for multiple organoids. While organoids are complex models of human organs, they are inherently less complex than the human organ but have proven particularly useful in drug discovery and modelling complex diseases including many cancers. Organoids are arguably more reliable than other forms of in vitro testing and it is likely that a multi-organoid simulation will eventually be found to be superior to small rodent and primate testing for preclinical drug development in terms of cost, time, and reliability. In the context of exploding AI, this vision of the future will be upon us very soon.

Near term development of the DeepNEU platform will focus on several areas. First, the simulation of the spinal cord and peripheral nervous system will need to be completed and integrated in the aiHumanoid. A working protype of the spinal cord is already in late development. In addition, other organoids like prostate, thymus, and parathyroid gland will need to be simulated and integrated. We are also introducing an altruism subsystem with the goal of maximizing the likelihood of producing predictions that are in the best interests of humanity. This feature is currently in testing and has not yet been integrated into the aiHumanoid simulation. Finally, some sensory inputs like sound and light are already part of the system but are quite rudimentary and need to be upgraded along with adding other forms of sensory inputs.

Our eventual goal is to replace pre-clinical wet lab, animal drug testing and eventually Phase 1 human trials with validated aiHumanoid simulations. Importantly, this ambitious goal appears to be consistent with the recent FDA decision to relax the requirements for animal testing before initiating human drug trials [138].

## Appendix A

### Organoid Markers

<u>Cardiac organoids</u> - Markers: TNNT2, ACTC1, MYH6, MYH7, NKX2.5

Reference: Richards, D. J., Coyle, R. C., Tan, Y., Jia, J., Wong, K., Toomer, K., … Mei, Y. (2017). Inspiration from heart development: Biomimetic development of functional human cardiac organoids. Biomaterials, 142, 112-123. https://doi.org/10.1016/j.biomaterials.2017.07.021

<u>Vascular organoids</u> - Markers: CD31, VE-Cadherin, VWF, SMA, NG2

Reference: Wimmer, R. A., Leopoldi, A., Aichinger, M., Wick, N., Hantusch, B., Novatchkova, M., … Penninger, J. M. (2019). Human blood vessel organoids as a model of diabetic vasculopathy. Nature, 565(7740), 505-510. https://doi.org/10.1038/s41586-018-0858-8

<u>Brain organoids</u> - Markers: SOX2, PAX6, TBR1, FOXG1, CTIP2

Reference: Lancaster, M. A. et al. (2013). Cerebral organoids model human brain development and microcephaly. Nature, 501(7467), 373-379.

<u>Cerebellar organoids?</u>

Purkinje cells - Markers: CALB1 (calbindin), PCP2 (Purkinje cell protein 2), L7

Granule cells - Markers: NEUROD1, PROX1, PAX6

Bergmann glia - Markers: GFAP (glial fibrillary acidic protein), S100B, BLBP (brain lipid-binding protein)

Molecular layer interneurons - Markers: PVALB (parvalbumin), CALB2 (calretinin)

GABAergic interneurons - Markers: GAD65/67 (glutamic acid decarboxylase)

Unipolar brush cells - Markers: MET (hepatocyte growth factor receptor), PLCB4 (phospholipase C beta 4)

Reference: Nayler, S. P., Powell, J. E., Vanichkina, D. P., Korn, O., Wells, C. A., Kanjhan, R., … Wolvetang, E. J. (2017). Human iPSC-Derived Cerebellar Neurons from a Patient with Ataxia-Telangiectasia Reveal Disrupted Gene Regulatory Networks. Frontiers in Cellular Neuroscience, 11, 321. https://doi.org/10.3389/fncel.2017.00321

<u>Brainstem organoids</u>

Neurons - Markers: MAP2, TUBB3, NEUN

Astrocytes - Markers: GFAP, S100B, ALDH1L1

Oligodendrocytes - Markers: OLIG2, MBP, SOX10

Serotonergic neurons (present in brainstem) - Markers: TPH2, SERT, VMAT2

Noradrenergic neurons (present in brainstem) - Markers: DBH, NET, TH

Cholinergic neurons (present in brainstem) - Markers: CHAT, VAChT, CHT1

Reference: Rifes, P., Isaksson, M., Rathore, G. S., Aldrin-Kirk, P., Møller, O. K., Barzaghi, G., … Kirkeby, A. (2020). Publisher Correction: Modeling neural tube development by differentiation of human embryonic stem cells in a microfluidic WNT gradient. Nature Biotechnology, 38(11), 1357. https://doi.org/10.1038/s41587-020-0590-4. Erratum for: Nature Biotechnology, 2020 Nov;38(11), 1265-1273.

<u>Endocrine organoids</u>

Pancreatic organoids - Markers: PDX1, SOX9, NKX6.1, INS, GCG

Reference: Hohwieler M, Illing A, Hermann PC, et al. Human pluripotent stem cell-derived acinar/ductal organoids generate human pancreas upon orthotopic transplantation and allow disease modelling. Gut. 2017 Mar;66(3):473-486. DOI: 10.1136/gutjnl-2016-312423. PMID: 27633923; PMCID: PMC5534761.

Adrenal organoids - Markers: CYP11B1, CYP11B2, CYP17A1, STAR, NR5A1

References: Ahn, C. H., Na, H. Y., Park, S. Y., Yu, H. W., Kim, S. J., Choi, J. Y., … Kim, J. H. (2022). Expression of CYP11B1 and CYP11B2 in adrenal adenoma correlates with clinical characteristics of primary aldosteronism. Clinical Endocrinology (Oxford), 96(1), 30-39. https://doi.org/10.1111/cen.14628.
Faienza, M. F., Chiarito, M., Baldinotti, F., Canale, D., Savino, C., Paradies, G., … Bertelloni, S. (2019). NR5A1 Gene Variants: Variable Phenotypes, New Variants, Different Outcomes. Sex Development, 13(5-6), 258-263. https://doi.org/10.1159/000507411.

Thyroid organoids - Markers: TTF1, PAX8, TPO, TG, NIS

Reference: Antonica, F., Kasprzyk, D. F., Opitz, R., Iacovino, M., Liao, X. H., Dumitrescu, A. M., … Costagliola, S. (2012). Generation of functional thyroid from embryonic stem cells. Nature, 491(7422), 66-71. https://doi.org/10.1038/nature11525.

### Gender Specific organoids

Mammary gland organoids - Markers: EPCAM, ER, PR, HER2, CK14

Reference: Sachs, N., et al. (2018). A living biobank of breast cancer organoids captures disease heterogeneity. Cell, 172(1-2), 373-386.e10.

Ovarian organoids - Markers: PAX8, WT1, FOXL2, ER, PR

Reference: Kopper, O., et al. (2019). An organoid platform for ovarian cancer captures intra- and interpatient heterogeneity. Nature Medicine, 25(5), 838-849.

Testicular organoids - Markers: VASA, GATA4, SOX9, AMH, PLZF

Reference: Alves-Lopes, J. P., Söder, O., & Stukenborg, J. B. (2017). Testicular organoid generation by a novel in vitro three-layer gradient system. Biomaterials, 130, 76-89.

### Internal organoids

Intestinal organoids - Markers: LGR5, CD44, EPHB2, OLFM4, ASCL2

Reference: Sato, T., et al. (2009). Single Lgr5 stem cells build crypt-villus structures in vitro without a mesenchymal niche. Nature, 459(7244), 262-265.

Liver organoids - Markers: ALB, AFP, CYP3A4, CK19, CK7

Reference: Huch M, et al. (2013) In vitro expansion of single Lgr5+ liver stem cells induced by Wnt-driven regeneration. Nature 494(7436):247-50.

Gallbladder organoids - Markers: KRT7, KRT19, CFTR, MUC1, MUC5AC

Reference: Koike, H., Iwasawa, K., Ouchi, R., et al. (2021). Engineering human hepato-biliary-pancreatic organoids from pluripotent stem cells. Nature Protocols, 16, 919-936. https://doi.org/10.1038/s41596-020-00441-w.

Kidney organoids - Markers: WT1, LHX1, PAX2, ECAD, SIX2

Reference: Takasato, M., Er, P. X., Chiu, H. S., Maier, B., Baillie, G. J., Ferguson, C., … Little, M. H. (2015). Kidney organoids from human iPS cells contain multiple lineages and model human nephrogenesis. Nature, 526(7574), 564-568. https://doi.org/10.1038/nature15695.

### Other organoids

Lung organoids - Markers: NKX2.1, SOX9, FOXJ1, MUC5AC, P63

Reference: Dye, B. R., Hill, D. R., Ferguson, M. A., Tsai, Y. H., Nagy, M. S., Dyal, R., … Spence, J. R. (2015). In vitro generation of human pluripotent stem cell derived lung organoids. Elife, 4, e05098. https://doi.org/10.7554/eLife.05098.

Skin organoids - Markers: KRT5, KRT14, IVL, LOR, P63

Reference: Lee, J., Rabbani, C. C., Gao, H., Steinhart, M. R., Woodruff, B. M., Pflum, Z. E., … Koehler, K. R. (2020). Hair-bearing human skin generated entirely from pluripotent stem cells. Nature, 582(7812), 399-404. https://doi.org/10.1038/s41586-020-2352-3.

### Organoid Primer

Reference: Zhao, Z., Chen, X., Dowbaj, A.M. et al. Organoids. Nat Rev Methods Primers 2, 94 (2022). https://doi.org/10.1038/s43586-022-00174-y

